# diplo-locus: A lightweight toolkit for inference and simulation of time-series genetic data under general diploid selection

**DOI:** 10.1101/2023.10.12.562101

**Authors:** Xiaoheng Cheng, Matthias Steinrücken

## Abstract

Whole-genome time-series allele frequency data are becoming more prevalent as ancient DNA (aDNA) sequences and data from evolve-and-resequence (E&R) experiments are generated at a rapid pace. Such data presents unprecedented opportunities to elucidate the dynamics of genetic variation under selection. However, despite many methods to infer parameters of selection models from allele frequency trajectories available in the literature, few provide user-friendly implementations for large-scale empirical applications. Here, we present `diplo-locus`, an open-source Python package that provides functionality to simulate and perform inference from time-series data under the Wright-Fisher diffusion with general diploid selection. The package includes Python modules as well as command-line tools and is available at: https://github.com/steinrue/diplo_locus.

## 1 Introduction

With the rapid advancements in sequencing technologies, large datasets of temporally stratified population genomic samples are increasingly common. On the one hand, improved techniques for processing ancient DNA (aDNA), that is, DNA from deceased remains, enable the collection of aDNA sequencing data for many samples in humans and other species. On the other hand, for evolution experiments with organisms in the laboratory, it has become increasingly affordable to sequence the genomes of samples taken at multiple time points, bringing forth a fast-growing body of evolve-and-resequence (E&R) datasets. These datasets enable observing changes in the genetic composition of a population across multiple time points, allowing detailed inference about the underlying processes.

One of the most intriguing forces that shape the dynamics of genetic variants is natural or artificial selection, as it reflects how the population adapts to its environment. The Wright-Fisher model and its diffusion approximation are commonly used to model the dynamics of allele frequencies over time, especially when selection is acting, and they have been applied to analyze temporal genetic data (*e*.*g*., Bollback et al., 2008; Malaspinas et al., 2012; Mathieson and McVean, 2013; Steinrücken et al., 2014; Steinrücken et al., 2016; He et al., 2020). Nevertheless, despite several existing approaches to infer selection parameters, most analyze the data exclusively under a model of additive (semi-dominant) selection, and are often applied to time-series at specific loci, rather than genome-wide. However, some recent approaches do leverage full genomic data (Mathieson and Terhorst, 2022; Whitehouse and Schrider, 2023).

Here, we present our open-source software package `diplo-locus` that implements the single-locus Wright-Fisher diffusion with piece-wise constant general diploid selection in a panmictic population (see Section S1 in the *Supplemental Material*). It can be used to simulate and analyze temporally sampled genetic data under a Hidden Markov Model (HMM) framework (Bollback et al., 2008). Using efficient Gaussian approximations of the allele frequency dynamics allows for likelihood-based inference of general diploid selection coefficients. With most of the computation vectorized, `diplo-locus` is able to analyze data at a large number of loci simultaneously without compromising efficiency and speed. We believe this lightweight toolkit is a useful addition to the suite of methods for the broader scientific community working with temporal genetic samples.

## 2 Methods

Our software package `diplo-locus` implements the Wright-Fisher diffusion and the HMM framework (see Section S1 in the *Supplemental Material*) in `Python3`. It requires the `NumPy` (Harris et al., 2020), `SciPy`, `pandas`, and `matplotlib` packages. Once installed, the command-line interface (CLI), `DiploLocus`, can be used to perform analyses in a Unix/Linux environment, or, alternatively, the respective functions can be imported from the `diplo locus` module in Python. Both the CLI implementation of this program and the functions of the module can perform likelihood computation and simulation with a wide range of customizable options. A detailed manual and tutorials can be found in the GitHub repository at: https://github.com/steinrue/diplo_locus.

### 2.1 Command-Line Interface DiploLocus (CLI)

To compute log-likelihoods, the command `DiploLocus likelihood` reads the genotype data either from a VCF file with a matching sample information file or from a table of allele counts with sampling times. The user can use command-line arguments to designate subsets of loci and filter loci by pooled minor allele frequency. For input as VCF in particular, the user can specify the subset of samples to be considered. If sampled individuals with diploid and individuals with pseudo-haploid genotypes should be analyzed together, the program accepts VCF files that contain both. To resolve ambiguities, the command-line options `--force dip` can be used to explicitly list a subset of individuals whose genotype should be interpreted as full diploid genotypes, whereas `--force hap` can be used to specify a subset of individuals whose genotype in the VCF file should be interpreted as pseudo-haploid: `0/0` corresponds to one reference allele, `1/1` corresponds to one alternative allele, and `0/1` is not permitted.

In addition to the input files, the user must provide an effective population size (`--Ne`), pergeneration recurrent mutation rate(s) (`--u[01|10]`), and the initial allele frequencies (`--initCond`), either as a fixed value or a distribution. For values of per-generation selection coefficients, the program accommodates two fitness parametrizations at a bi-allelic locus with alleles a and A (see Table S1 in the *Supplemental Material*): By relative fitness of heterozygotes *s*_aA_ and homozygotes *s*_AA_, or by selection *s* = *s*_AA_ and dominance coefficients *h*, with *s*_aA_ = *h · s*. The program provides options to compute log-likelihoods on a linear or geometric grid of values for each selection parameter or at a fixed value. The program also allows for selection models with selection coefficients changing over time in a piece-wise constant manner.

Besides computing log-likelihoods on a specified grid, the program can interpolate 1-dimensional log-likelihood surfaces and output on-and off-grid maximum likelihood estimates (MLEs) of the selection coefficients at each locus for increased precision. Furthermore, the program offers the choice to output *p*-values computed from the likelihood ratio statistic using a *χ*^2^ distribution with 1 degree of freedom (Self and Liang, 1987).

To simulate temporal samples and allele frequency trajectories, the command `DiploLocus simulate` allows users to simulate an arbitrary number of independent replicates under specified population and selection parameters. The tool requires a constant effective population size and constant mutation rates. In addition, it can simulate using a model of constant selection throughout time, as well as piece-wise constant selection coefficients. Multiple filtering options based on sample or population frequencies are also provided. As output, the program produces sample allele counts at the specified sampling times, and, optionally, their population allele frequency trajectories. Lastly, the program also allows for convenient visualization of the sample and population frequency trajectories for select replicates.

### 2.2 Python module diplo locus

The module `diplo_locus.likelihood` contains the class `SelHmm`, which initializes the HMM for computation using specified population and selection parameters as described in Section 2.1. The function `SelHmm.computeLikelihood()` then computes per-locus log-likelihoods based on these parameters for a given number of replicates of temporal samples. The module `diplo locus.simulate` contains the function `simulateSamples()` to simulate independent replicates based on given population and selection parameters. It returns replicate samples with the corresponding underlying trajectories. Both, analysis and simulation can accommodate piece-wise constant temporal changes in selective pressure.

## 3 Results

### 3.1 Statistical Performance

We tested the statistical performance of `diplo-locus` on data simulated using either `diplo-locus`, see Figure 1 and Section S2 in the *Supplemental Material*, or `SLiM4` (Haller and Messer, 2023), see Section S3. We observe from the ROC curves presented in Figure 1a and Figure 1c, that the likelihood ratio of the estimated MLEs divided by neutrality (*s* = 0) computed using `diplo-locus` can distinguish replicates simulated under selection from neutral replicates. Figure 1b demonstrats that the MLEs of *s*_*AA*_ capture the true value used for the simulations accurately, and Figure 1d shows that `diplo-locus` can estimate the time of selection onset *t*_*o*_ accurately for strong selection, if this time is not known a priori. Additional simulation results in Section S2 and Section S3 of the *Supplemental Material* further demonstrate the power and accuracy of `diplo-locus`.

**Figure 1.**
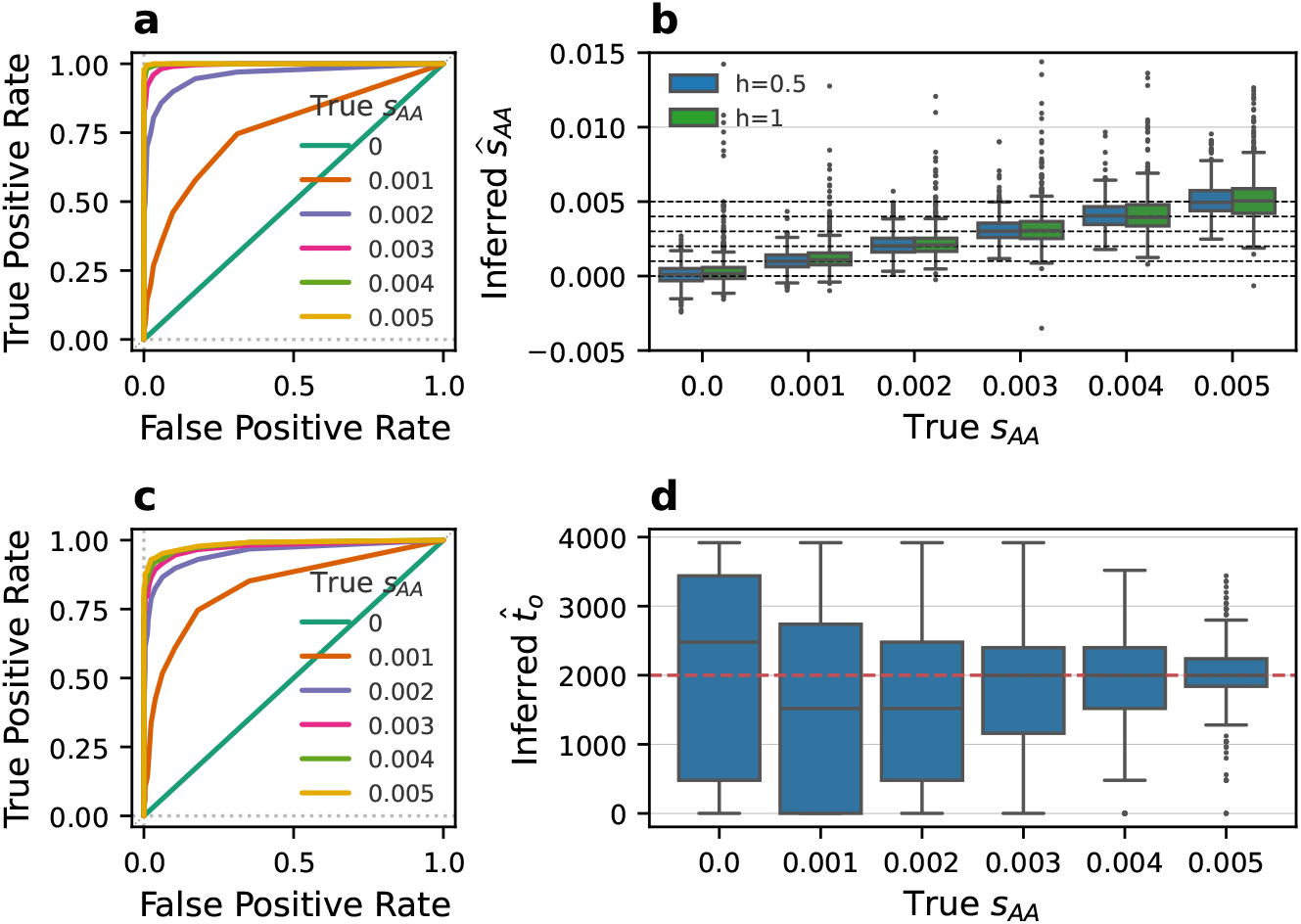
Receiver operating characteristic (ROC) curves demonstrate power to distinguish replicates simulated under **a)** additive selection and **c)** selection with dominance *h* = 1 from neutral replicates using likelihood ratio tests. Curves based on replicates of 9 temporal samples of 20 diploid individuals, each 500 generations apart, and the diploid population size is *N*_*e*_ = 10^4^. **b)** Boxplot of MLEs ŝ_AA_ for the same set of simulated data used in a) & c). **d)** Marginal point estimates of the time of selection onset *t*_*o*_. The selection coefficient was set to 0 before *t*_*o*_ = 2, 000, and to *s*_*AA*_ afterwards. Here, 50 diploid individuals were sampled at each time.)

**Figure 2.**
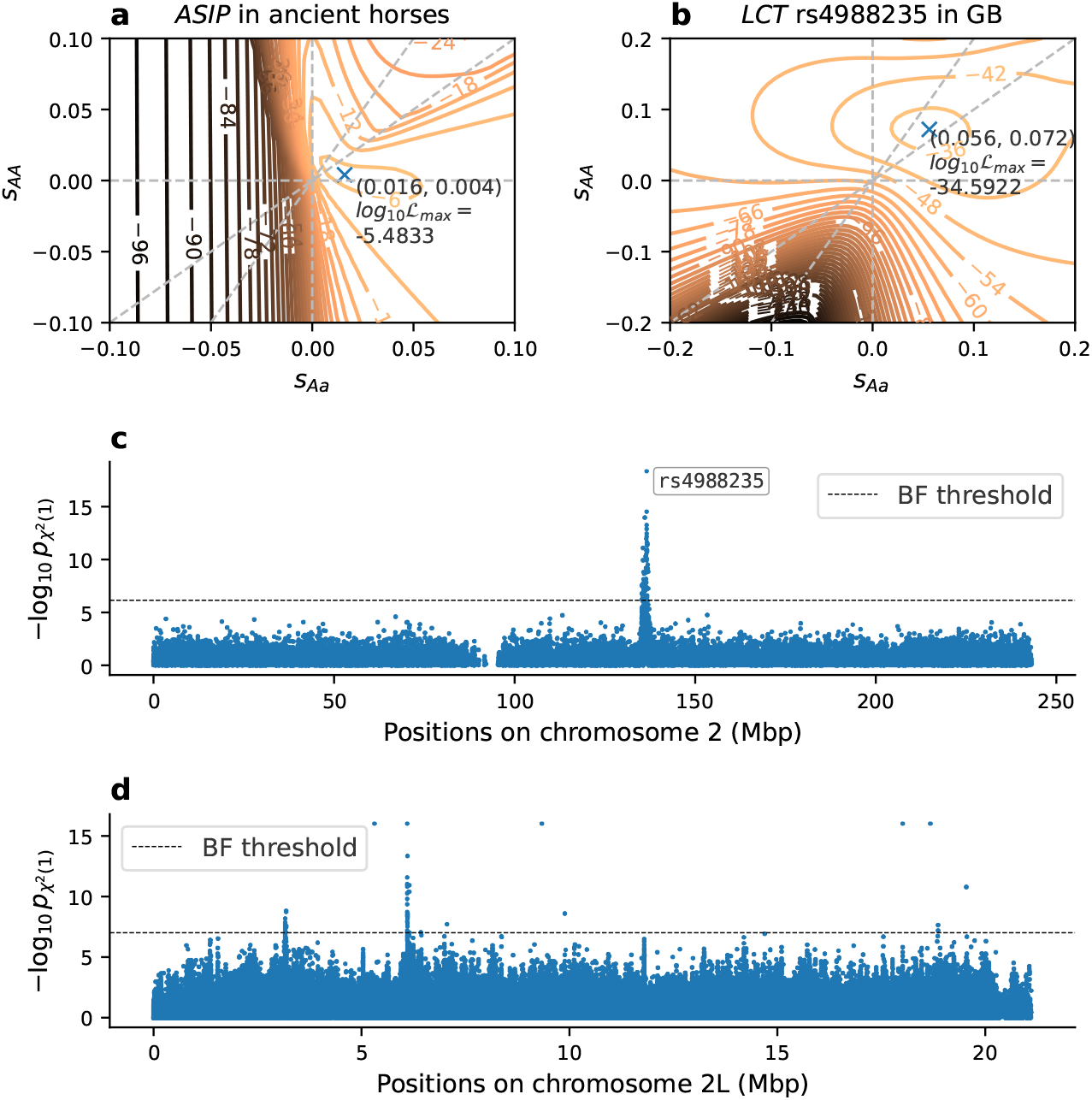
Log-likelihood surface across two-dimensional grid of diploid selection coefficients *s*_Aa_ and *s*_AA_, computed on 101*×*101 linear grid: **a)** *ASIP* locus in ancient horses from Ludwig et al. (2009) and **b)** SNP `rs4988235` in the *LCT* /*MCM6* locus in ancient human samples from Great Britain (GB). Diagonal dashed gray lines indicate full dominance (*s*_Aa_ = *s*_AA_) and additivity (*s*_Aa_ = *s*_AA_*/*2). **c)** Manhattan plot of *p*-values for filtered SNPs on Chromosome 2 in the GB aDNA human data. **d)** Manhattan plot of *p*-values for filtered SNPs on Chromosome 2L from *D. simulans* E&R experiment, combined across 10 biological replicates using Fisher’s method.

### 3.2 Computational Performance

We compared the performance of `diplo-locus` against several other methods to infer selection coefficients from temporal data presented in the literature: `LLS` (Taus et al., 2017), `WFABC` (Foll et al., 2015), and `bmws` (Mathieson and Terhorst, 2022). We simulated 12,000 replicates of temporal data in a variety of scenarios of different length: 160 generations or 4000 generations; additive (*h* = 0.5) or dominance (*h* = 1) selection; and constant selection or time varying selection, see Section S4 of the *Supplemental Material* for a detailed description. Table 1 shows the runtimes for the different methods to analyze all simulated replicates in each scenario, using a single core of an Intel Ice Lake processor (2.9GHz) with 32GB RAM. Note that the methods from the literature can only be applied to a subset of the scenarios, and thus do not have runtimes reported for all. In addition, Figure S14 and Figure S15 in the *Supplemental Material* show the accuracy of the estimates for the respective methods.

**Table 1.** Comparison of runtimes for different methods. # Gener. is number of generations simulated, *h* indicates dominance coefficient, and “Var. *s*” indicates whether *s* varies over time. Runtimes are given as hours required to analyze all 12,000 replicates in each scenario. We ran `WFABC` and `bmws` for 24 hours and extrapolated the runtimes based on number of completed replicates. Missing value indicates that the method cannot analyze data in the respective scenario

| # Gener. | $h$ | Var. $s$ | diplo-locus | LLS | WFABC | bmws |
| --- | --- | --- | --- | --- | --- | --- |
| 160 | 0.5 | No | 0.054 | 0.006 | 296.952 | 162.193 |
| 160 | 0.5 | Yes | 0.050 | - | - | 90.831 |
| 160 | 1.0 | No | 0.048 | 0.046 | 2055.296 | - |
| 160 | 1.0 | Yes | 0.047 | - | - | - |
| 4000 | 0.5 | No | 0.077 | 0.006 | 691.143 | 357.558 |
| 4000 | 0.5 | Yes | 0.068 | - | - | 379.343 |
| 4000 | 1.0 | No | 0.068 | 0.938 | 248.255 | - |
| 4000 | 1.0 | Yes | 0.068 | - | - | - |

In the scenarios with additive selection, the method `LLS` essentially performs logistic regression, which is highly optimized, and thus the fastest. In all other scenarios, `diplo-locus` is faster or equally fast compared to `LLS`. Both `diplo-locus` and `LLS` are substantially faster than `WFABC` and `bmws`. Due to the vectorized implementation, `diplo-locus` shares the transition matrices for the underlying HMM across replicates, resulting in substantial computational speed-up. Furthermore,

Figure S14 and Figure S15 in the *Supplemental Material* show that `diplo-locus` is at least as accurate as the most accurate method among the other three in all scenarios. These results demonstrate that `diplo-locus` is an efficient and flexible method that produces accurate estimates of general diploid selection coefficients in a wide range of scenarios.

### 3.3 Application to Empirical Data

To showcase the capabilities of `diplo-locus`, we applied the method to three empirical datasets: Temporal data for pigmentation alleles at the ASIP locus in ancient horses (Ludwig et al., 2009), see Section S5.1 in the *Supplemental Material* ; Chromosome 2 of ancient humans from Great Britain (GB), dated more recent than 4,500 years before present, extracted from the Allen Ancient DNA Resource (Mallick et al., 2024, v54.1), see Section S5.2; and E&R data for Chromosome 2L of *Drosophila simulans* exposed to a new temperature regime for 60 generations (Barghi et al., 2019), see Section S5.3.

Figure 1a shows the likelihood surface for general diploid selection at the ASIP locus, which is consistent with previous findings of balancing selection (Steinrücken et al., 2014). In Figure S16 in the *Supplemental Material*, we show likelihood surfaces for three different times of origin of the selected allele. We find that the likelihood is maximized for a time of origin immediately before the selected allele was observed in the sample, and the corresponding surface is shown in Figure 1a. Figure 1b shows the likelihood surface for the well-characterized SNP `rs4988235` on Chromosome 2 in the GB aDNA human dataset, which regulates expression of the lactase *LCT* gene (Troelsen et al., 2003), with a point MLE of (*s*_Aa_, *s*_AA_) = (0.056, 0.072). Figure 1c shows a Manhattan plot of the *p*-values under additive selection for all SNPs on Chromosome 2 in the GB human aDNA dataset, and the SNP `rs4988235` is the most significant signal. Figure 1d shows a Manhattan plot of the *p*-values under additive selection for Chromosome 2L of the *D. simulans* E&R data, combined across 10 biological replicates using Fisher’s method. Figure S20 and Figure S21 in the *Supplemental Material* show detailed plots of the annotated genes in the vicinity of the two clusters of significant *p*-values. The genes in these regions are related to transmembrane transporter activity and kinase signaling. A detailed analysis of genome-wide results for these datasets is beyond the scope of this manuscript and will be addressed in future work.

We note that `diplo-locus` assumes that the temporal samples originate from a single well-mixing population of constant size. Thus, for the analysis, it is important to specify the effective population size accurately. For the analyses presented here, we provide details in the respective sections of the *Supplemental Material*, but in brief, we use the effective size estimated by Der Sarkissian et al. (2015) for the ASIP data in horses, the harmonic mean of the sizes estimated by Browning et al. (2015) for the human data from Great Britain, and the size estimated by Barghi et al. (2019) for the *Drosophila* data.

As a benchmark, using an Intel Ice Lake processor (2.9 GHz), it takes about 4 hours to compute log-likelihoods for the 520 samples of the GB dataset at the 69,903 genotyped SNPs (after filtering) across a 51-point grid of *s*_AA_ values. Note that, because the likelihood values are stored in memory before writing output or maximizing, the program might consume more memory than the user’s system capacity in cases of a large number of loci or dense parameter grids. Thus, to analyze a larger number of loci, it might be necessary to split the analysis into several parallel batches. As a guideline: The above analysis of 69,903 genotyped SNPs across a 51-point grid of *s*_AA_ required 4 GB to run, and the required memory scales linearly with number of SNPs and number of gridpoints.

## 4 Discussion

We present a toolkit developed in Python for analyzing temporal genetic samples under the Wright-Fisher model, with a focus on inferring general diploid selection. Our software can compute likelihoods for time series data, perform likelihood-based inference, and simulate replicates of such data. We assessed the efficiency and accuracy of the inference using our software and showcased the versatility of the CLI `DiploLocus` and the python module `diplo locus`. We believe this package will be a valuable addition to the toolkit of the population genetics community.

## Web Resources

Our software `diplo-locus` is available at: https://github.com/steinrue/diplo_locus. Scripts to recreate the simulations and analyses presented in this manuscript are available at: https://github.com/steinrue/diplo_locus_manuscript_figs.

## Data Availability Statement

The temporal data for the pigmentation alleles in ancient horses is given by Ludwig et al. (2009). The ancient human data is extracted from the Allen Ancient DNA Resource v54.1 described by Mallick et al. (2024), available at: https://doi.org/10.7910/DVN/FFIDCW (Dataverse V7). The E&R data for *D. simulans* is described by Barghi et al. (2019), available at: https://doi.org/10.5061/dryad.rr137kn.

## Acknowledgements

We thank Adam Fine, Jeremy Berg, John Novembre, Maanansa Raghavan, Constanza de la Fuente Castro, and Dylan Sosa for helpful discussions and feedback on this project.

