## Supplementary Material for "diplo-locus: A lightweight toolkit for inference and simulation of time-series genetic data under general diploid selection"

### S1 Hidden Markov Model to Compute Likelihoods of Temporal Samples

#### S1.1 Diploid Wright-Fisher Model and Temporal Samples

We assume a panmictic diploid population of effect size $N_e$ and consider a bi-allelic locus with alleles a and A. Denote the per-site per-generation mutation probability for mutations a $\rightarrow$ A by $u_{\mathrm{aA}}$ and for mutations A $\rightarrow$ a by $u_{\mathrm{Aa}}$. Denote the random population allele frequency of the focal allele (without loss of generality A) in generation $t \geq 0$ by $X_t \in [0, 1]$. In addition, assume that the alleles at the respective locus are evolving under general diploid selection. That is, the relative fitness of an AA homozygote, an aA heterozygote, and an aa homozygote is $1 + s_{\mathrm{AA}}$, $1 + s_{\mathrm{aA}}$, and 1, respectively, for selection coefficients $s_{\mathrm{AA}}$ and $s_{\mathrm{aA}}$ (Table S1). Alternatively, this model can be parameterized using a parameter for the strength of selection $s$, and a dominance coefficient $h$, such that $s_{\mathrm{AA}} = s$, and $s_{\mathrm{aA}} = h \cdot s$.

Table S1: Fitness schemes for diploid genotypes. Here "A" is the focal allele. Parameters $s$ and $h$ denote the selection coefficient and dominance coefficient, respectively.

| Genotype | AA | Aa | aa |
|---|---|---|---|
| Fitness | $1 + s_{\mathrm{AA}}$ | $1 + s_{\mathrm{aA}}$ | 1 |
| | $1 + s$ | $1 + h \cdot s$ | 1 |

Suppose $\mathbf{n} = (n_1, ..., n_K)$ alleles are sampled from the population at generation $t_k \geq 0$, $k \in \{1, \ldots, K\}$, with $K$ being the number of sampling time points. Within each time-stratified pool of sampled alleles, denote the numbers of focal alleles by $\mathbf{d} = (d_1, ..., d_K)$. To compute the likelihood of this temporal data, we employ a Hidden Markov Model (HMM) framework (Bollback et al., 2008; Mathieson and McVean, 2013; Steinrücken et al., 2014), where the population allele frequency $X_t$ is the hidden state, evolving according to the Wright-Fisher model, and the temporal data is the emission, where the number of samples with the focal alleles at generation $t_k$ is binomially distributed conditional on the population allele frequency $X_{t_k}$.

To facilitate the numerical computations in this HMM framework, we discretize the continuous hidden state space $[0, 1]$ of $X_t$ into a finite number of $M+1$ intervals. To this end, we choose $M+1$ equidistant points $f_m := \frac{m}{M} \in [0, 1]$ with $m \in \{0, \ldots, M\}$, as representative for each interval. Note that $f_0 = 0$, and $f_M = 1$. Using these representative points, we can define interval boundaries $g_\ell := \frac{1}{2}(f_{\ell-1} + f_\ell)$ for $\ell \in \{1, \ldots, M\}$, $g_0 := -\infty$, and $g_{M+1} := \infty$, which in turn can be used to define intervals $I_m := (g_m, g_{m+1})$. Note that $f_m \in I_m$ holds for all $m \in \{0, \ldots, M\}$. With this discretization, we define the random variables $F_t \in \{f_0, \ldots, f_M\}$ for $t \geq 0$ that represent the allele frequency dynamics of $X_t$ in the discretized state space such that $\{X_t \in I_m\} \approx \{F_t = f_m\}$.

##### S1.1.1 Initial Distribution

At generation $t = 0$, denote the probability density function for the distribution of the initial population allele frequency $X_0$ by $\pi(x)$. Here, we consider several initial distributions: a fixed frequency, uniform distribution, or Beta distribution, which is the stationary distribution for the recurrent mutation model (eg. Durrett, 2008, Ex. 7.17). With $x \in [0, 1]$, these are respectively given by

$$\pi^{\delta}(x) = \delta_y(x),$$
$$\pi^{\text{unif}}(x) = \mathbf{1}_{[0,1]}(x),$$
and
$$\pi^{\text{beta}}(x) = \frac{1}{B(\alpha, \beta)}(1-x)^{\alpha-1}x^{\beta-1},$$

with $\alpha = 4N_e u_{\text{Aa}}$, $\beta = 4N_e u_{\text{aA}}$, and the Beta function $B(\alpha, \beta) = \frac{\Gamma(\alpha)\Gamma(\beta)}{\Gamma(\alpha+\beta)}$, where $\Gamma(y)$ denotes the Gamma function. Here $\delta_y$ denotes the dirac delta at $y \in (0,1)$, and $\mathbf{1}_A(x)$ denotes the indicator function, which is 1 if $x \in A$, and 0 otherwise. Note that for $\alpha = \beta = 1$, the Beta distribution corresponds to the uniform distribution. Lastly, the initial distribution for the discretized frequency $F_0$ is then given by

$$\begin{aligned} \mathbb{P}_\Theta\{F_0 = f_m\} &:= \mathbb{P}_\Theta\{X_0 \in I_m\} \\ &= \int_{x=g_m \vee 0}^{g_{m+1} \wedge 1} \pi(x)\, dx, \end{aligned} \tag{S1.1}$$

where $\pi(\cdot)$ is the respective initial distribution and $\Theta$ is the collection of all model parameters.

##### S1.1.2 Transition

Under the Wright-Fisher model, assuming the coefficients for both mutation and selection are small, the increment of the allele frequency from generation $t-1$ to the next generation $t$ can be approximated by a normal distribution as

$$(X_t - X_{t-1}) \sim \mathcal{N}\Big(\mu(X_{t-1}), \sigma^2(X_{t-1})\Big),$$

where $\mathcal{N}(\mu, \sigma^2)$ denotes a normal distribution with mean $\mu$, variance $\sigma^2$, and

$$\begin{cases} \mu(x) & = \big[u_{\text{aA}}(1-x) - u_{\text{Aa}}x\big] + x(1-x)\big[s_{\text{aA}}(1-2x) + s_{\text{AA}}x\big] \\ \\ \sigma^2(x) & = \frac{1}{2N_e} \cdot x(1-x), \end{cases}$$

see, for example, Chapter 7 by Durrett (2008). Our implementation of the model allows for piece-wise constant changes in the selection coefficient, but we omit this from the notation for clarity.

For convenience, denote by $\varphi(x; \mu, \sigma^2)$ the probability density for a normally distributed random variable with mean $\mu$ and variance $\sigma^2$. Then, we define the probability of transitioning from the discretized allele frequency $F_{t-1}$ at generation $t-1$ to the discretized allele frequency $F_t$ in generation $t$ as

$$\begin{aligned} &\mathbb{P}_\Theta\{F_t = f_m | F_{t-1} = f_{m'}\} \\ &\quad := \mathbb{P}_\Theta\{X_t \in I_m | X_{t-1} = f_{m'}\} \\ &\quad = \int_{x=g_m}^{g_{m+1}} \varphi\big(x; \mu_\Theta(f_{m'}), \sigma^2_\Theta(f_{m'})\big)\, dx. \end{aligned} \tag{S1.2}$$

Note that our choice of $g_0 := -\infty$ and $g_{M+1} := \infty$ assigns the mass of the normal distribution outside of the interval $[0, 1]$ to the respective boundary points.

Furthermore, note that in the discretized model, the probability of transitioning across multiple generations can be computed using matrix exponentiation. That is, for $t' < t$, we have

$$\mathbb{P}\{F_t = f_m | F_{t'} = f_{m'}\} = \left(A^{(t-t')}\right)_{m',m}, \tag{S1.3}$$

where $A$ is the $(M + 1) \times (M + 1)$ one-step transition matrix with coefficients $A_{i,j} := \mathbb{P}\{F_t = f_j | F_{t-1} = f_i\}$.

##### S1.1.3 Emission

Conditional on the population frequency of the focal allele $X_{t_k}$ at a time $t_k$ when a sample is taken, the probability of sampling $d_k$ focal alleles out of $n_k$ alleles sampled in total is binomially distributed. Thus, denoting the number of sampled focal alleles by $Y_{t_k}^{(n_k)}$, we have

$$\mathbb{P}\{Y_{t_k}^{(n_k)} = d_k | X_{t_k} = x\} = \binom{n_k}{d_k} x^{d_k} (1-x)^{n_k - d_k}.$$

Again, using the discretized hidden state space to facilitate computation in the HMM, we can define the emission probability as

$$\begin{aligned} &\mathbb{P}\{Y_{t_k}^{(n_k)} = d_k | F_{t_k} = f_m\} \\ &\quad := \mathbb{P}\{Y_{t_k}^{(n_k)} = d_k | X_{t_k} = f_m\} \\ &\quad = \binom{n_k}{d_k} (f_m)^{d_k} (1 - f_m)^{n_k - d_k}. \end{aligned} \tag{S1.4}$$

#### S1.2 Likelihood of Temporal Samples

The likelihood of the observed temporal data under the HMM with the discretized hidden state space can be computed using the standard forward algorithm (eg. Bishop, 2016, Ch. 13.2.2) by iteratively computing the forward variable

$$a_t(m) := \mathbb{P}_\Theta\{F_t = f_m, Y_{t_\ell}^{n_\ell} = d_\ell \;\forall \ell \text{ s.t. } t_\ell \leq t\}, \tag{S1.5}$$

for all $t_k$. To this end, we initialize the forward variable as

$$a_0(m) = \mathbb{P}_\Theta\{F_0 = f_m\} \times \begin{cases} \mathbb{P}\{Y_0^{(n_1)} = d_1 | F_0 = f_m\}, & \text{if } t_1 = 0, \\ 1, & \text{otherwise,} \end{cases}$$

where the initial probability is defined in equation (S1.1) and the emission probability in equation (S1.4). Define $t_0 := 0$ for convenience. If the forward variable is computed for a given $t_{k-1}$, we can iteratively compute it for $t_k$ using

$$a_{t_k}(m) = \left(\sum_{m'=0}^{M} \mathbb{P}_\Theta\{F_{t_k} = f_m | F_{t_{k-1}} = f_{m'}\} \, a_{t_{k-1}}(m')\right) \mathbb{P}\{Y_{t_k}^{(n_k)} = d_k | F_{t_k} = f_m\}$$

where the transition probability is defined in equation (S1.2) and (S1.3), and the emission probability in equation (S1.4). Lastly, after computing the forward variable $a_{t_k}(m)$ iteratively for all $t_k$ with $k \in \{0, \ldots, K\}$, thus up to and including the time of the last sample, we can compute the likelihood of the data by summing over all hidden states at the last time point:

$$\ell_\Theta(\mathbf{d}, \mathbf{n}) := \mathbb{P}_\Theta\{Y_{t_\ell}^{n_\ell} = d_\ell \,\forall 1 \leq \ell \leq K\} = \sum_{m=0}^{M} a_{t_K}(m).$$

Note that in the implementation of `diplo-locus`, we use scaling factors (Bishop, 2016, Ch. 13.2.4) to avoid underflow in the floating point arithmetic. That is, rather than computing the forward variable $a_{t_k}(m)$ defined in equation (S1.5), we compute the normalized version

$$\hat{a}_{t_k}(m) := \mathbb{P}_\Theta\{F_{t_k} = f_m | Y_{t_\ell}^{n_\ell} = d_\ell \,\forall \ell \leq k\},$$

by renormalizing the forward variable at each step of the iteration, which also yields the scaling factors

$$c_{t_k} = \mathbb{P}_\Theta\{Y_{t_k}^{n_k} = d_k | Y_{t_\ell}^{n_\ell} = d_\ell \,\forall \ell < k\}.$$

These can then in turn be used to compute the likelihood of the data using

$$\ell_\Theta(\mathbf{d}, \mathbf{n}) = \mathbb{P}_\Theta\{Y_{t_\ell}^{n_\ell} = d_\ell \,\forall 1 \leq \ell \leq K\} = \prod_{k=1}^{K} c_{t_k}.$$

#### S1.3 Maximum Likelihood estimates of selection coefficients and Likelihood Ratios

Here, we are primarily interested in estimating the parameters of the diploid selection model. To emphasize this, we define $\Theta(s_{\mathrm{aA}}, s_{\mathrm{AA}})$ as the set where all other model parameters are fixed and only the parameters for the diploid selection vary. Alternatively, we can also define $\Theta(s)$ for models of selection where only one parameter varies: for example, for additive selection, $s_{\mathrm{aA}} = \frac{1}{2}s$ and $s_{\mathrm{AA}} = s$. We furthermore define $\Theta_0 := \Theta(0, 0)$ as set of parameters with selection coefficients equal to zero, thus neutrality.

To obtain the Maximum Likelihood Estimate (MLE) of the selection coefficients, we can maximize the likelihood using

$$(\widehat{s_{\mathrm{aA}}}, \widehat{s_{\mathrm{AA}}}) := \underset{(s_{\mathrm{aA}}, s_{\mathrm{AA}})}{\operatorname{argmax}} \, \ell_{\Theta(s_{\mathrm{aA}}, s_{\mathrm{AA}})}(\mathbf{d}, \mathbf{n})$$

or

$$\widehat{s} := \underset{s}{\operatorname{argmax}} \, \ell_{\Theta(s)}(\mathbf{d}, \mathbf{n}).$$

The likelihood ratio for this MLE is then given by

$$\mathrm{LR} := \Big[\frac{\ell_{\Theta(\widehat{s})}(\mathbf{d}, \mathbf{n})}{\ell_{\Theta_0}(\mathbf{d}, \mathbf{n})}\Big]^2,$$

and the logarithm of the likelihood ratio is given by

$$\log \mathrm{LR} = 2\Big(\log \ell_{\Theta(\widehat{s})}(\mathbf{d}, \mathbf{n}) - \log \ell_{\Theta_0}(\mathbf{d}, \mathbf{n})\Big). \tag{S1.6}$$

### S2 Inferring parameters from replicates simulated using `diplo-locus`

#### S2.1 Simulation set-up

Using the module `diplo_locus.simulate`, we simulated 500 replicates each for 6 values of the selection coefficient $s_{\mathrm{AA}} \in \{0, 0.001, 0.002, 0.003, 0.004, 0.005\}$ for the AA homozygote, with dominance coefficients $h \in \{0, 0.5, 1\}$, corresponding to recessive, additive, and dominant fitness, respectively. To investigate models of heterozygote advantage, we furthermore simulated replicates using the same set of selection coefficients, but for $s_{\mathrm{aA}}$, the selection coefficient of heterozygotes, either with dominance $h = 5$, and thus $s_{\mathrm{AA}} = \frac{1}{h} s_{\mathrm{aA}}$, or with $s_{\mathrm{AA}} = 0$, that is, perfect over-dominance. The selection pressure was set constant throughout the time period simulated.

We set the effective population size to $N_e = 10,000$, use symmetric per-site per-generation mutation rates $u_{aA} = u_{Aa} = 0$, and set the initial condition to be either a fixed frequency of 0.01, or following Watterson's neutral distribution (Watterson, 1975), mimicking standing variation, where

$$\begin{aligned} &\mathbb{P}\{i \text{ focal alleles among } 2N_e \text{ haploids}\} \\ &\quad = \mathbb{P}\big\{\text{population allele frequency at } \tfrac{i}{2N_e}\big\} \\ &\quad = \frac{1/i}{\sum_{k=1}^{2N_e}(1/k)}. \end{aligned} \tag{S2.7}$$

Note that the latter also represents the stationary distribution under the Poisson random field model (Sawyer and Hartl, 1992). We then sampled 40 haploids (20 diploids) at nine points in time, equidistant every 500 generations.

We condition the simulations on the selected allele not being lost. To this end, we removed replicates whose allele frequency dropped below $1/(4N_e)$ at any time throughout the simulation. To remove uninformative samples, we also filtered the simulated replicates by their pooled minor allele frequency (MAF). A replicate was removed if, after pooling all the samples, its MAF is lower than 0.05. After removing replicates, we re-simulated replacement replicates until the total number of valid replicates reached 500. Simulation scripts and the scripts to generate figures presented here can be found in the repository `https://github.com/steinrue/diplo_locus_manuscript_figs`.

#### S2.2 Likelihood computation

We used the class `SelHmm` in the module `diplo_locus.likelihood` to compute likelihoods for the simulated replicates. We computed log-likelihoods assuming either a uniform initial distribution or a fixed initial frequency of 0.01. The log-likelihoods were computed along a geometric grid of $s$ values in $[-0.75, 0.75]$ that is symmetric around 0 with 51 points. The selection coefficient is assumed to be constant during the time period considered. For the effective population size and the mutation rates, we fixed the values as used for the simulations.

Once the log-likelihoods were computed on the grid for each replicate, we used the function `scipy.interpolate.interp1d` to obtain the log-likelihoods as a function of $s$ in the 1-dimensional continuous space, and determine an off-grid maximum using `scipy.optimize`. We report the MLE of the selection coefficient, $\hat{s}_{\mathrm{MLE}}$, and its corresponding log-likelihood ratio (LLR) statistic, obtained by taking twice the difference between the log-likelihood at $s = \hat{s}_{\mathrm{MLE}}$ and that at $s = 0$, see equation (S1.6).

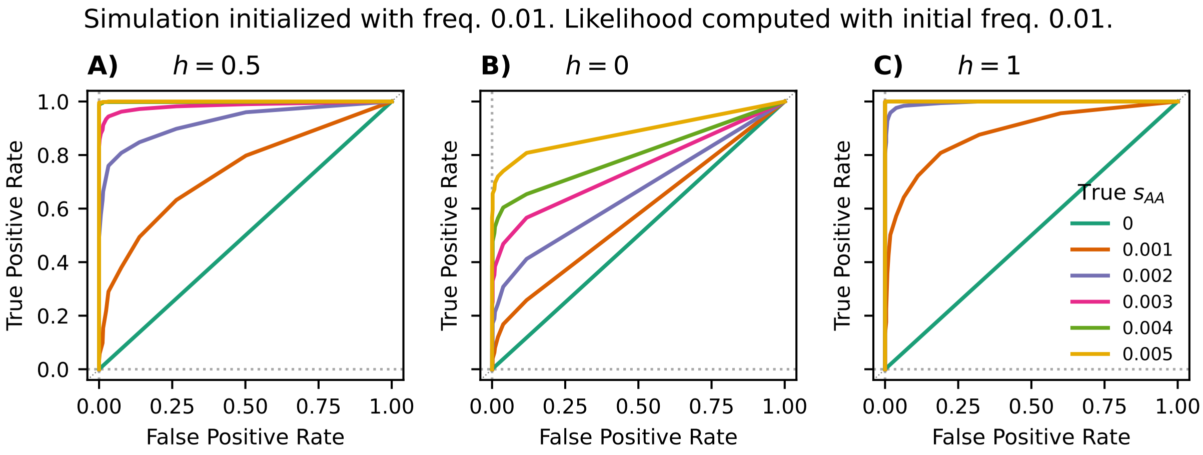

Figure S1: ROC curves for LLR computed using `diplo-locus` for samples simulated to start at a fixed frequency of 0.01. Likelihood computation assumes a fixed starting frequency 0.01 and fixed dominance coefficient of A) $h = 0.5$ (additive), B) $h = 0$ (recessive), and C) $h = 1$ (dominant).

#### S2.3 Power to distinguish selection from neutrality

The LLR is well-powered to distinguish sites evolving under selection above a certain strength from sites evolving under neutrality in the simulations, exhibited in the Receiver operating characteristic (ROC) curves in Figure S1, S2, S3. Among the three dominance coefficients examined, the statistical power of the method is lowest for $h = 0$ and highest for $h = 1$. This is conceivable, since under a model of dominance, the initial frequency increase of alleles starting at low frequency is strongest, and thus more replicates show a discernible signature.

For simulations initiated at a fixed frequency of 0.01 (Figure S1 and S2), the initial condition used by the likelihood computation does not substantially change the statistical power. It is possible that the influence of the initial condition is stronger if fewer individuals are sampled at fewer time points. However, if the initial allele frequency in a particular application is unknown, we recommend using the uniform distribution, since it is agnostic to any specific assumptions and performs rather favorably.

For simulations initiated with Watterson's neutral distribution, we observe higher true positive rate (TPR) for $s = 0.001$, 0.002, and 0.003 for $h = 0.5$ or 0 (Figure S3) compared with those starting from a fixed frequency of 0.01 (Figure S2). This behavior can be explained by the mean difference in the initial frequencies. Under Watterson's neutral distribution with $N_e = 10^4$, the starting allele frequency is on average higher than 0.01, resulting in stronger frequency changes, and thus a stronger signature of selection detectable by the method. Meanwhile, the TPR for stronger selection ($s_{\mathrm{AA}} \geq 0.002$ for $h = 1$) does not approach 1 as quickly (Figure S3C), which is likely due to the removal of replicates by the MAF filter that start at a high frequency and fix immediately.

In addition to ROC curves based on LLR statistics, we also computed p-values for the simulated replicates. Because in the different scenarios the null hypothesis ($s_{\mathrm{AA}} = 0$) is embedded in the one-dimensional space of alternative hypothesis ($s_{\mathrm{AA}} \neq 0$), the likelihood ratio statistic given in equation (S1.6) asymptotically follows a $\chi^2$ distribution (Self and Liang, 1987) with one degree

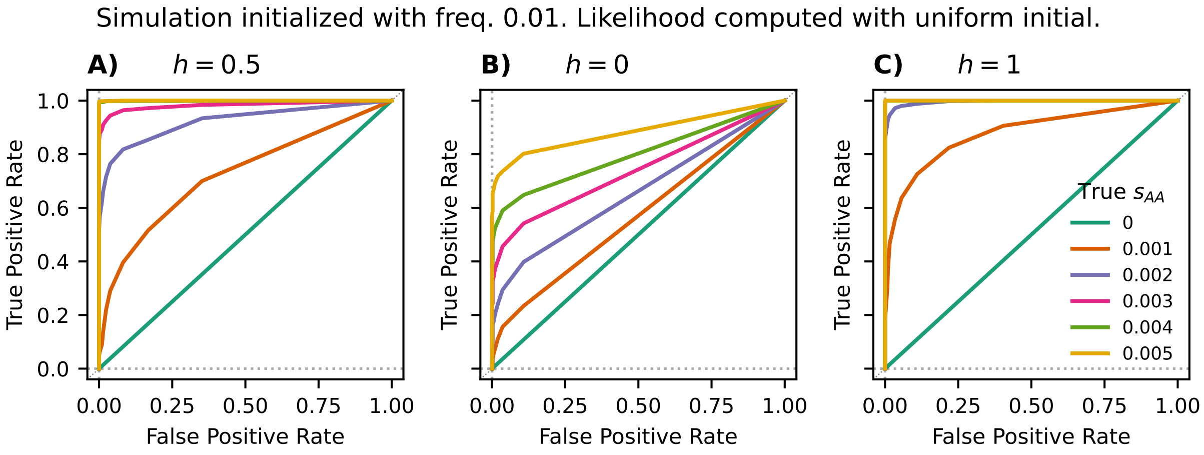

Figure S2: Receiver operating characteristic (ROC) curves for the likelihood ratio (LLR) computed using `diplo-locus` for samples simulated to start at a fixed frequency of 0.01. Likelihood computation assumes a uniform initial frequency distribution and uses a fixed dominance coefficient of A) $h = 0.5$ (additive), B) $h = 0$ (recessive), and C) $h = 1$ (dominant).

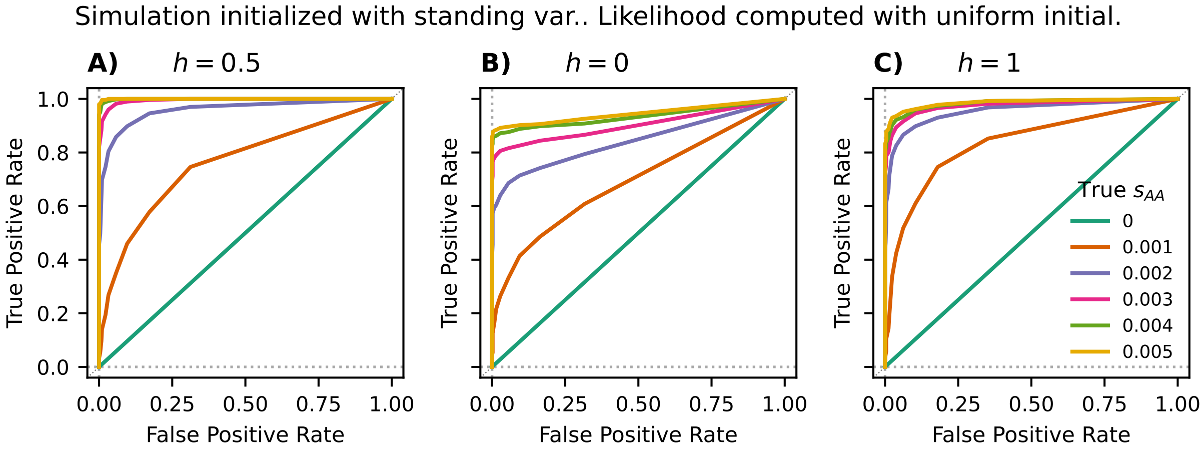

Figure S3: ROC curves for the LLR computed using `diplo-locus` for samples simulated to start at a frequency drawn from Watterson's neutral distribution (see equation (S2.7)). Likelihood computation assumes uniform initial conditions and fixed dominance coefficient of A) $h = 0.5$ (additive), B) $h = 0$ (recessive), and C) $h = 1$ (dominant).

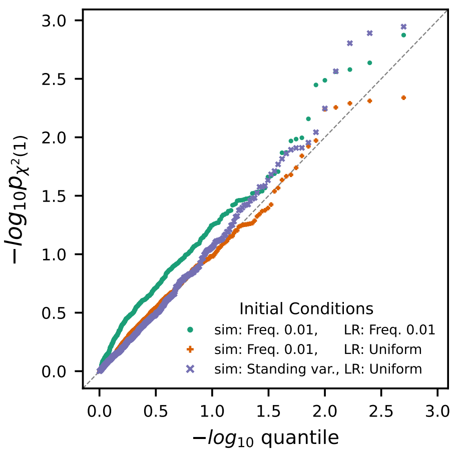

Figure S4: Quantile-quantile (QQ) plot of the p-values computed using the asymptotic $\chi^2(1)$ from neutral simulations in the case of additive selection ($h = 0.5$) with the respective initial conditions for simulation and likelihood computation.

of freedom. Figure S4 demonstrates, for additive selection ($h = 0.5$), that the p-values computed using a $\chi^2(1)$ distribution for the neutral replicates are well calibrated, and thus the asymptotic result can be used for the finite sampling scheme.

##### S2.4 Accuracy in estimating selection coefficients

Next, we examined the accuracy of parameter estimation based on the computed likelihood. For the selection coefficient of homozygotes, $s_{\text{AA}}$, the inferred MLEs are close to the true values, which are always covered by the inter-quartile range (body of each box in Figure S5, S6 and S7). In the recessive case, under the uniform distribution (recommended if the initial frequency is unknown), the MLEs exhibit a tighter distribution than if the initial distribution for computing likelihoods matches the simulated initial frequency of 0.01. Similar to their performance in detection power, the difference in the distributions of the MLEs is negligible for most cases.

The sampling scheme in a particular application does not always capture enough information about the allele frequency dynamics to allow reliable inference, regardless of the inference framework. In the simulations presented here, we sample the population every 500 generations. Under this setting, for an allele that rises from a low frequency to fixation in fewer than 500 generations, the sampled data would only allow to infer that a strongly selected allele quickly rose in frequency. However, accurate inference of the respective selection coefficient is only possible if the rise in frequency occurs over several sampled time points. The converse is also true: If the sampled time points are close together, changes in allele frequency due to weak selection might not be detectable. This poses a practical limit on the range of selection parameters that can be characterized using the given dataset and sampling scheme, of which the user should be mindful. In specific situations that a user is interested in, we thus strongly recommend simulation studies to characterize power and accuracy in the specific scenario.

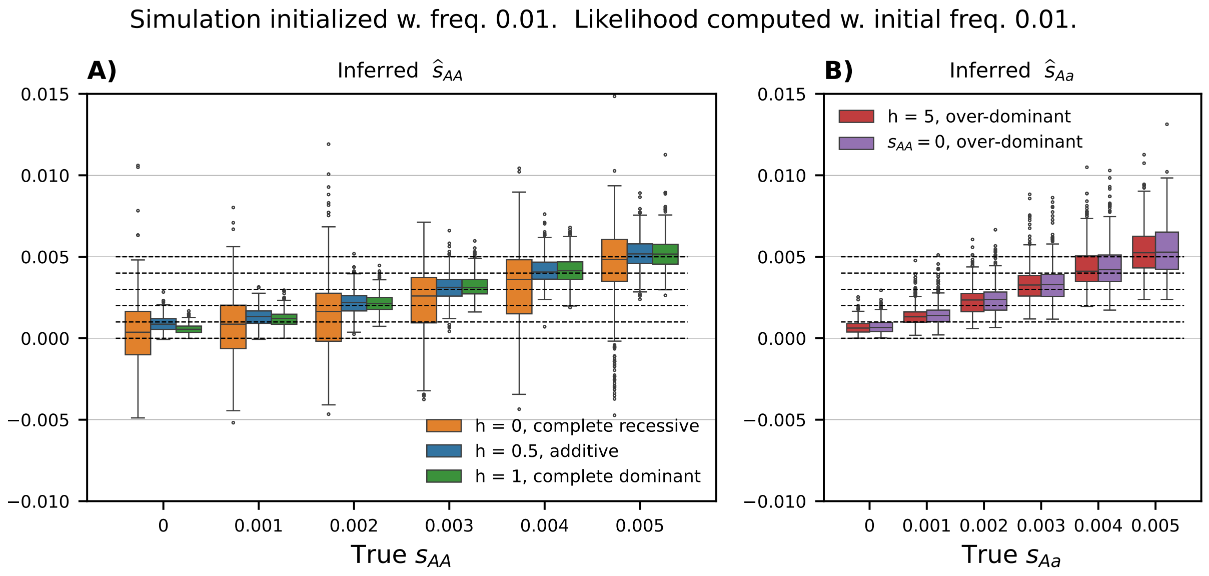

Figure S5: Distributions of MLEs of the selection coefficient A) $\hat{s}_{AA}$ with varying dominance or B) $\hat{s}_{Aa}$ with heterozygote advantage (over-dominance), inferred using a fixed initial frequency of 0.01. Replicates were simulated with an initial condition at a fixed frequency of 0.01.

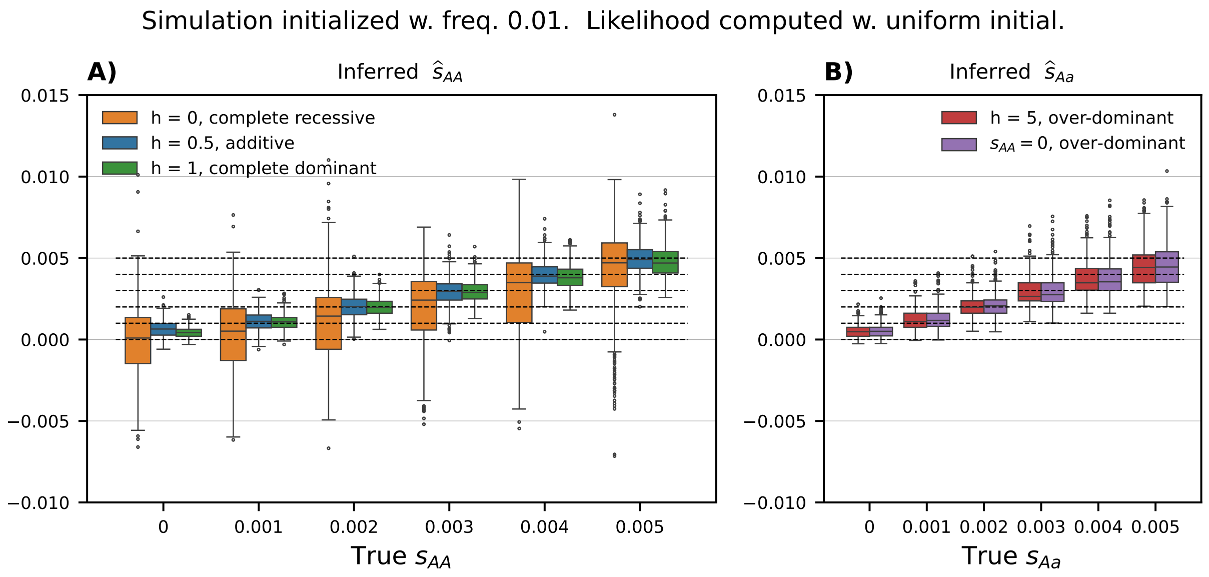

Figure S6: Distributions of MLEs of the selection coefficient A) $\hat{s}_{AA}$ with varying dominance or B) $\hat{s}_{Aa}$ with heterozygote advantage (over-dominance), inferred under a uniform initial condition. Replicates were simulated with an initial condition at a fixed frequency of 0.01.

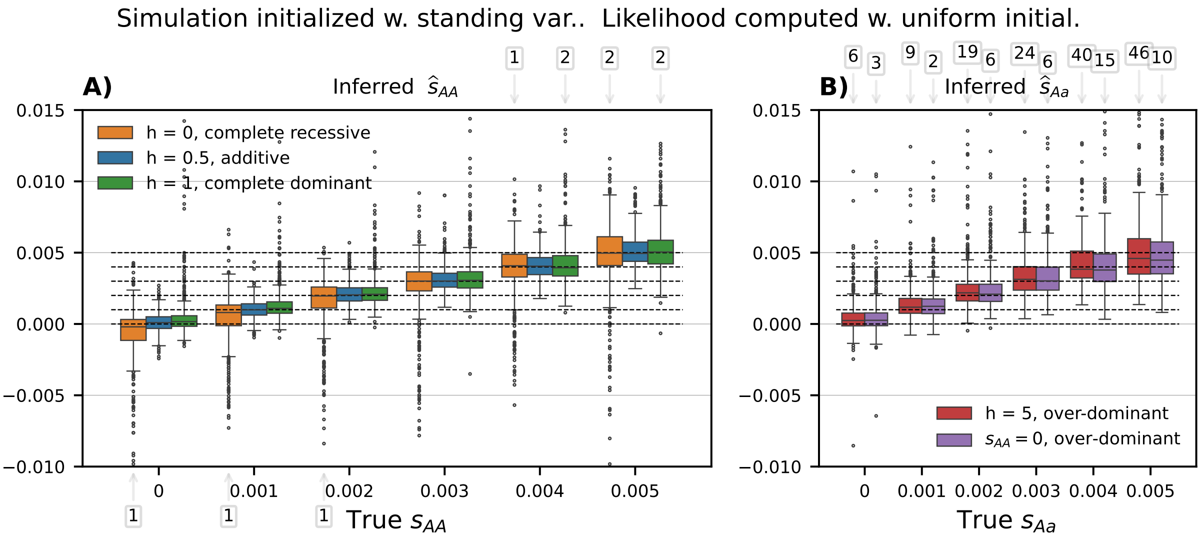

Figure S7: Distributions of MLEs of the selection coefficient A) $\hat{s}_{AA}$ with varying dominance or B) $\hat{s}_{Aa}$ with heterozygote advantage (over-dominance), inferred under a uniform initial condition. Replicates were simulated with initial frequency drawn from Watterson's neutral distribution. Numbers in boxes indicate number of replicates outside of plotting range.

##### S2.5 Inferring the onset of selection

In addition to scenarios where selection is constant throughout the time samples are taken, we investigated applying `diplo-locus` to estimate the time when selection begins. To this end, similar to Section S2.1, we simulated temporal samples of a selected allele from a population of size $N_e = 10,000$ with mutation rates $u_{aA} = u_{Aa} = 0$ that evolved for 4,000 generations. The initial frequency of the focal allele was set to 0.01 for each simulated replicate, and again conditioned on the population frequency never falling below $1/(4N_e)$ and a pooled MAF of 0.05 in the sample. We simulated under additive selection with coefficients $s_{\mathrm{AA}} \in \{0, 0.001, 0.002, 0.003, 0.004, 0.005\}$. For each coefficient we simulated 500 replicates with constant selection for all 4,000 generations and 500 replicates, where the selection coefficient was set to 0 for the first 2,000 generations, and then set to $s_{\mathrm{AA}}$ for generation 2,000 to 4,000. For each replicate, we sampled 100 haploid samples at 9 equidistantly spaced timepoints 500 generations apart.

We then computed likelihoods for all simulated replicates with `diplo-locus`, using $N_e = 10,000$ and $u_{aA} = u_{Aa} = 0$ as simulated, and the uniform distribution for the initial condition. For each replicate, we computed these likelihoods on a 2 dimensional grid $(t_o, s_{AA})$: For each pair of 51 possible times of onset $t_o$, equidistantly spaced between 0 and 4,000 generations, and 71 additive selection coefficients $s_{AA}$ on a geometric grid between -0.1 and 0.1, including 0. We then chose the point $(\hat{t_o}, s_{\hat{A}A})$ on this grid that maximized the likelihood as the MLE for the time of the onset of selection $t_o$ and the selection coefficient $s_{AA}$. We also computed likelihoods under constant selection for all 4,000 genrations. To asses significance, we computed the log-likelihood ratio (LLR) statistic as the difference between the log-likelihood at the MLE and the log-likelihood for $s_{AA} = 0$, and multiplied this difference by two.

Figure S8A) shows a QQ-plot of the ranked p-values computed from the replicates simulated under neutrality against the ranked p-values expected under neutrality. Here, we compute the p-values using two different approaches: Asymptotically, when performing a likelihood ratio test where the null hypothesis is nested in the parameter space of alternative hypotheses, the likelihood-ratio statistic is distributed according to a $\chi^2$-distribution, where the degree of freedom is equal to the difference in dimension of the parameter space for the alternative hypothesis and the null hypothesis (Self and Liang, 1987). The null hypothesis of neutrality $s_{AA} = 0$ is nested in the parameter space of alternative hypotheses for inferred time of onset $t_o$ and selection coefficient $s_{AA}$, and the difference in dimension is equal to two. We thus compute p-values under a $\chi^2$-distribution with degree of freedom 1 and 2, and indeed observe that using degree of freedom 2 yields well calibrated p-values.

In addition, Figure S8C) depicts receiver operator characteristic (ROC) curves when comparing the LLR statistics of replicates simulated with onset of selection against the statistics computed on neutral simulations. Figure S8B) and Figure S8D) show the corresponding marginal estimates of the additive selection coefficient $s_{AA}$ and the time of onset $t_o$, respectively. We observe limited power and inaccurate estimates when selection is weak, but for strong selection, power to detect non-neutral replicates is substantial, and the estimates are accurate. Our method `diplo-locus` is thus well suited to detect the onset of selection in scenarios where this time is not known a priori.

### S3 Inferring diploid selection from replicates simulated using `SLiM`

#### S3.1 `SLiM` simulation set-up

For the `SLiM` simulations, we used `msprime` v1.3.3 (Baumdicker et al., 2022) to initialize a novel selected allele on a neutral genomic background. Specifically, we used the “Discrete Time Wright-Fisher” (DTWF) mode of `msprime` to simulate 200 kbp genomic segments neutrally evolving along a constant-sized human-chimpanzee demographic history, as described by Cheng and DeGiorgio (2020), that is, two populations of size $N_e = 10,000$ that diverged from a common ancestral population of size $N_e = 10,000$ and evolve for 200,000 generations in isolation. We used the chimp outgroup to emulate misclassifying the ancestral allele. At generation 200,000, we use `SLiM` v4.2.2 (Haller and Messer, 2023) to identify the segregating site closest to the center of the simulated genomic region, and introduce single-locus diploid selection of the derived allele at this locus with the given parameters, which corresponds to selection on standing variation. We then output temporal genetic samples in the subsequent 4,000 generations. Two hundred replicates were generated for each selection scenario. For each replicate, nine samples of 40 haploids are taken every 500 generations during the 4,000 generations. Note that the mutation rate at the selected site and the sampling scheme here are identical to those in Section S2.1. Simulation scripts, `eidos` files, and the scripts to generate the figures presented here can be found in the repository `https://github.com/steinrue/diplo-locus_manuscript_figs`.

We applied the same conditioning to the simulated data as we applied to the datasets simulated using `diplo-locus`, see Section S2.1: If the allele was either lost or the pooled MAF of the sample was less than 0.05, we restarted the entire simulation pipeline to obtain a new replicate. For selection parameters in the simulation, we considered dominance coefficient $h \in \{0, 0.5, 1\}$ with selection coefficient $s_{\mathrm{AA}} \in \{0, 0.001, 0.002, 0.003, 0.004, 0.005\}$, as well as $h = 5$ with $s_{\mathrm{aA}} \in$

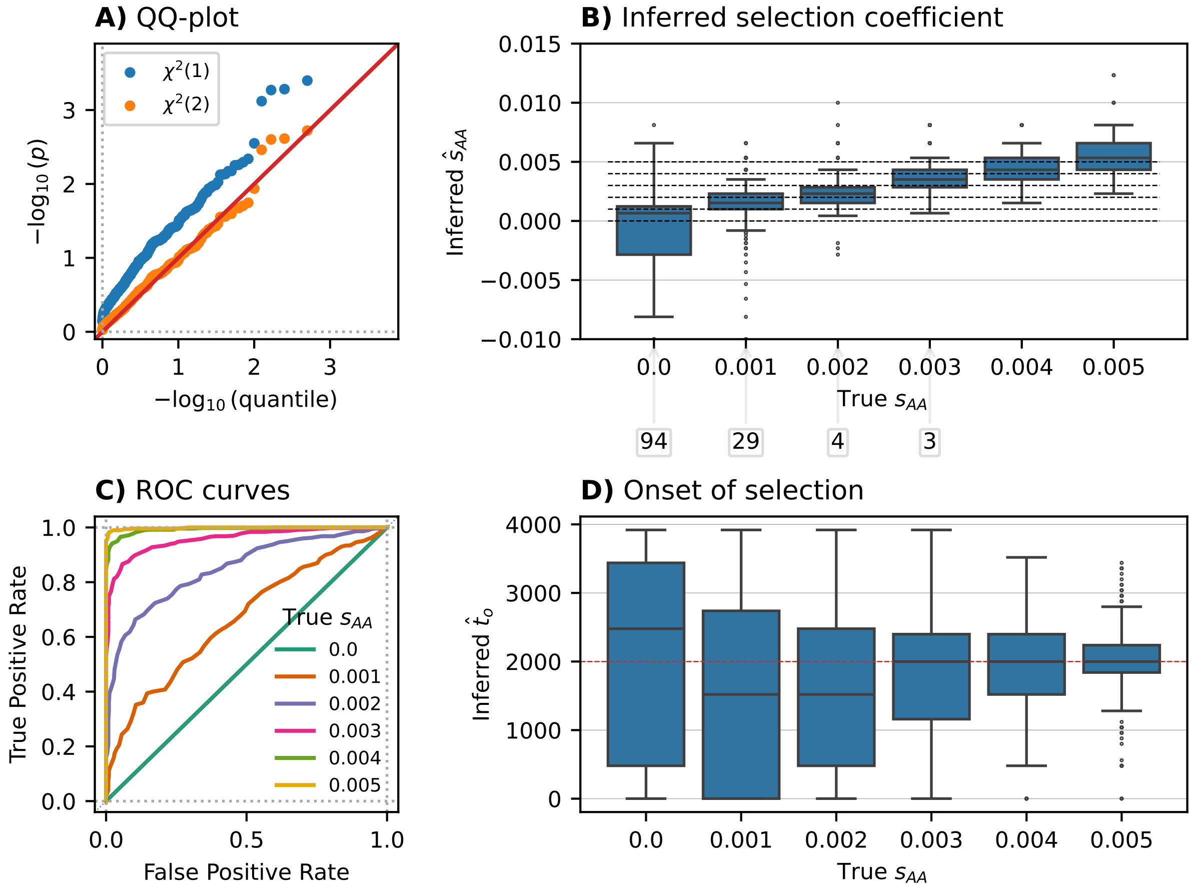

Figure S8: Point estimates and likelihood based testing when inferring onset of selection. A) QQ-plot of p-values for non-neutrality and inferred onset, computed using different asymptotics. B) Boxplot of marginal point estimates of selection coefficients $s_{AA}$. C) ROC curves indicating power under different simulated true selection coefficients $s_{AA}$. D) Boxplot of marginal point estimates of time of selection onset $t_o$. Plots based on 500 simulated replicates in each scenario.

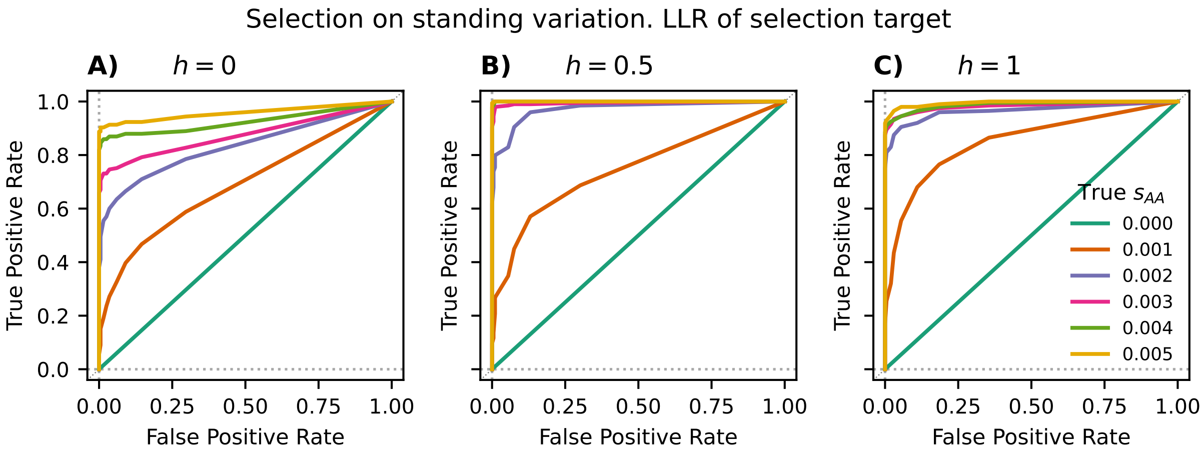

Figure S9: ROC curves for the LLR computed using `diplo-locus` for the selected mutation in samples simulated by `SLiM` after initialization using `msprime`. Likelihood computation assumes a uniform initial condition and fixed dominance coefficients A) $h = 0$, B) $h = 0.5$, or C) $h = 1$.

$\{0.001, 0.002, 0.003, 0.004, 0.005\}$.

We applied `diplo-locus` to the temporal data at the selected locus to compute the MLEs of the selection coefficients and likelihood ratio test statistics to reject neutrality using the same approach as detailed in Section S2.2. The results are presented in Figure S9 and Figure S10. As expected, these closely match the results obtained when simulating datasets from standing variation using `diplo-locus`, see Figure S3 and Figure S7.

#### S3.2 Identifying locus under selection and estimating selection coefficients

In addition to characterizing selection at a specific locus, we also explored how `diplo-locus` can be applied to identify the locus under selection in a given genomic region. To this end, we applied `DiploLocus` to compute LLRs and MLEs at all loci of the simulated 200 kbp regions (after applying the MAF filter). For each replicate, we choose the locus with the highest LLR within the 180 kbp region centered on the true selection target as the putative target of selection. In the neutral case ($s_{\mathrm{aA}} = 0$ or $s_{\mathrm{AA}} = 0$), we still report the distance from the distinguished locus where the selection coefficient was explicitly specified.

Figure S11 shows the distance of the estimated locus from the true target of selection, Figure S12 shows the corresponding ROC curves rejecting neutrality at this locus, and Figure S13 the corresponding accuracy of the MLEs for the selection coefficients at this locus. For the latter, we report the absolute value of the MLE rather than the signed value, because we cannot guarantee that the putative locus is the true target. At neutral sites linked to the target, either the ancestral or the derived alleles could be linked to the selected allele at the target site. For selection coefficients below 0.002, the distance to the real target shows high variability and the MLEs are biased. This is likely due to the fact genetic drift and randomness in the sampling lead to outliers among the neutral loci that mask weak selection at the target locus. However, for coefficients above 0.003, the accuracy increases and the bias vanishes. Thus, this procedure can be used to identify and

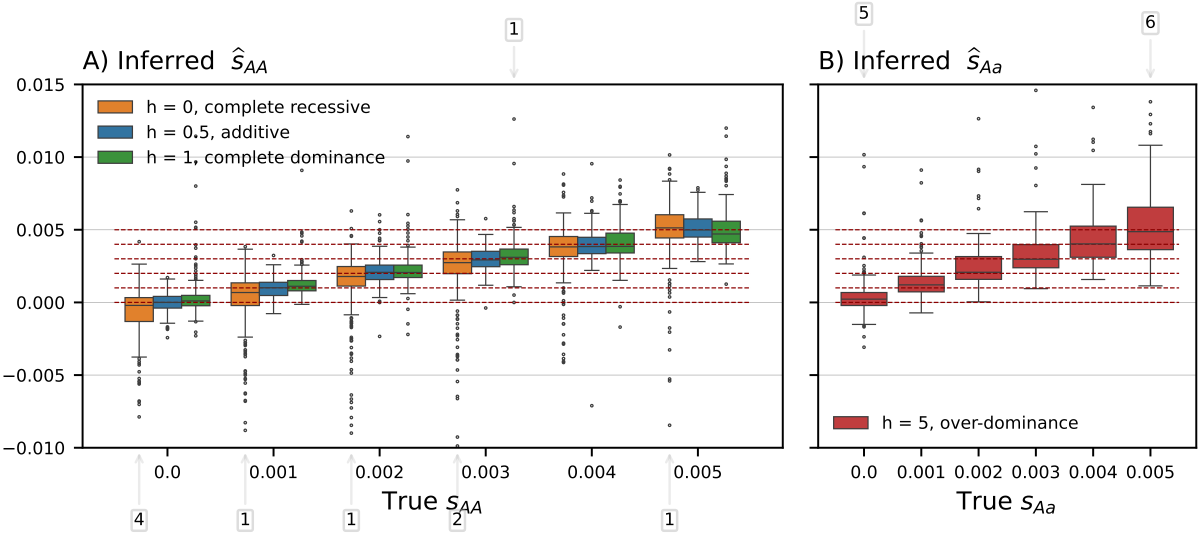

Figure S10: Distribution of MLEs of selection coefficient for the selected mutation, inferred under a uniform initial condition for sets of simulated loci with different dominance coefficients. Numbers in boxes indicate number of replicates outside of plotting range.

characterize targets of selection, if the selection is sufficiently strong.

#### S4 Comparison of methods to infer selection from time series data

Our method `diplo-locus` can be applied to temporal allele frequency data to estimate piecewise constant general diploid selection coefficients. We thus simulated several relevant scenarios and compared the performance of `diplo-locus` with methods from the literature that perform similar inference: `LLS` (Taus et al., 2017), `WFABC` (Foll et al., 2015), and `bmws` (Mathieson and Terhorst, 2022).

For each of our simulated scenarios, we chose the initial frequency from standing variation, as described in Section S2.1, and sampled 40 haploid samples at 9 timepoints. Since possible applications of the methods include Evolve & Resequence data, as well as ancient DNA, we simulated scenarios of different length: One set for 160 generations with $N_e = 1,000$ (sampling every 20 generations), and one set for 4000 generations with $N_e = 10,000$ (sampling every 500 generations). We simulated under additive selection ($h = 0.5$) and dominance ($h = 1$). Since the scenarios differ in length, selection has to be stronger in the shorter scenarios to be detectable. We thus used selection coefficients $s \in \{0, 0.01, 0.02, 0.03, 0.04, 0.05\}$ when simulating for 160 generations and selection coefficients $s \in \{0, 0.001, 0.002, 0.003, 0.004, 0.005\}$ when simulating for 4000 generations. We furthermore produced one set of simulations where the selection coefficients is constant for the entire duration, and another set of simulations where the selection coefficient is 0 for the first and the last quarter, and only non-zero in the middle two quarters. For each scenario and combinations of parameters, we simulated 2000 replicates, conditioning on the selected allele not being lost and an MAF of at least 0.05 when pooling all samples, as in Section S2 and Section S3.

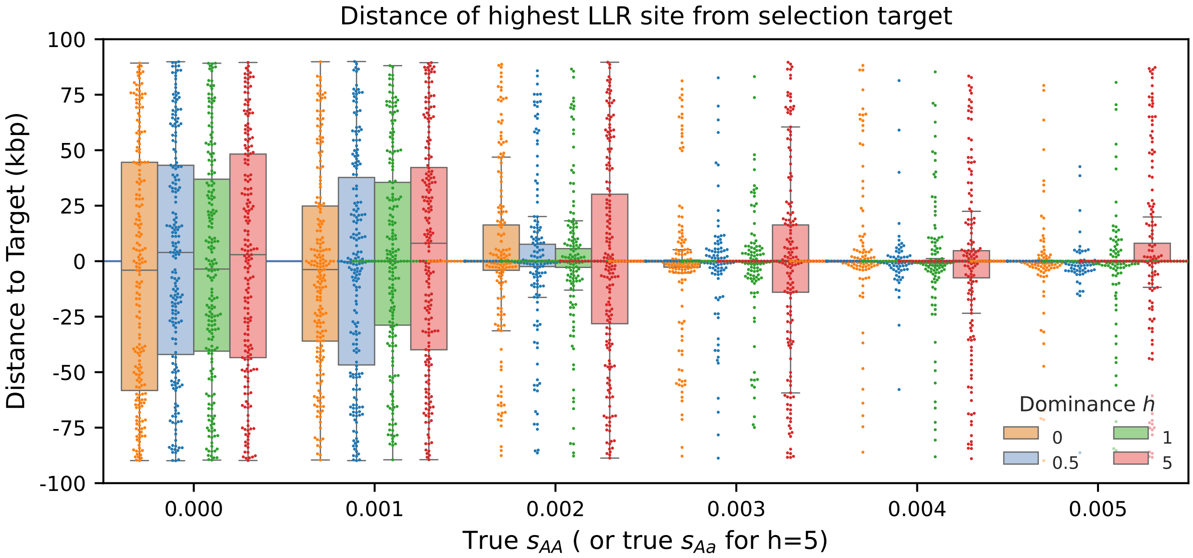

Figure S11: Distribution of the distance from the position with the highest LLR to the position of the true target of selection. Boxplots show the quartiles. For $s_{\mathrm{aA}} = 0$ or $s_{\mathrm{AA}} = 0$, the "true target" evolves neutrality, thus the distribution of the identified position is uniform in the genomic region.

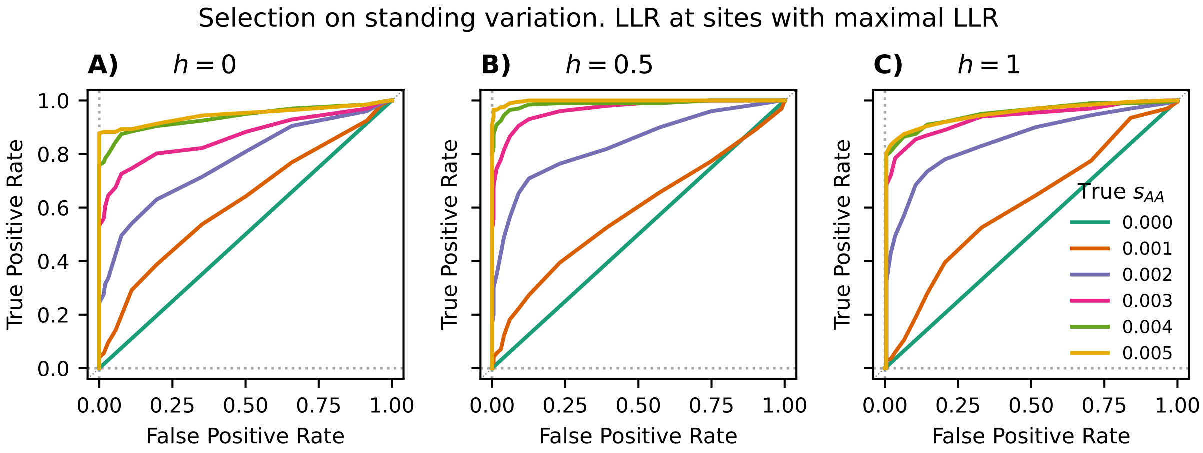

Figure S12: Receiver operating curves (ROCs) curves for the LLR computed using `diplo-locus` at the locus with highest LLR in simulated genomic region. Likelihood computation assumes a uniform initial condition and fixed dominance coefficients A) $h = 0$, B) $h = 0.5$, or C) $h = 1$.

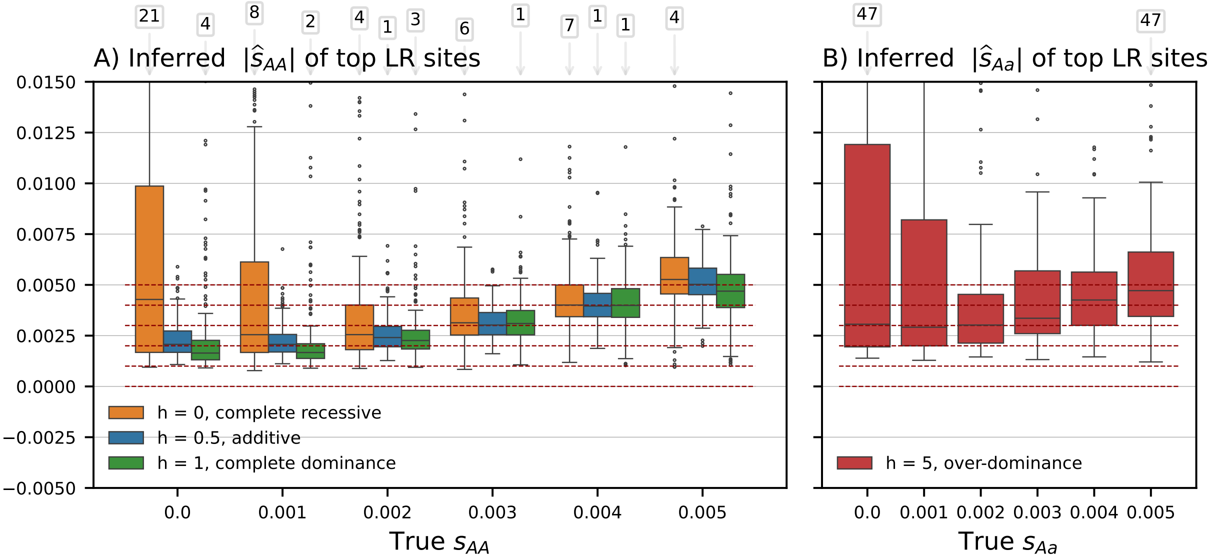

Figure S13: Distributions of absolute value of selection coefficient MLEs inferred at locus with highest LLR under a uniform initial condition. Numbers in boxes indicate replicates outside of plotting range.

We apply each of the methods in the respective scenario that they were designed for: All four methods can be applied to estimate a single constant additive selection coefficient. For the Bayesian method `WFABC`, we choose a Gaussian prior on s with mean 0 and standard deviation 0.075 when simulating for 160 generations and 0.0075 when simulating for 4000 generations. For `bmws`, we used a stronger regularization ($\lambda = 7$) in the constant coefficient case. In addition, the method was designed for additive selection and can thus not be applied in the scenarios with dominance. For `WFABC`, we set the prior on h such that $h = 1$ always holds to analyze the data simulated under dominance. `diplo-locus` and `bmws` are the only methods that can be applied in the scenarios where the selection coefficient changes over time. For `diplo-locus`, we fixed the times of change and specify that selection is 0 outside of the middle interval. `bmws` was not designed for an abrupt change in the selection coefficient as simulated here, but for more gradual changes. We thus ran the method allowing it to estimate changing selection coefficients for each generation (with the regularization recommended by the authors $\lambda = 4.5$), and then take the average of the coefficients estimated in the middle interval. Lastly, the only method that can estimate time-changing non-additive selection coefficients is `diplo-locus`.

We report the total runtime to analyze all simulated replicates in each scenario in Table 1 in the main text. Furthermore, Figure S14 and Figure S15 depict boxplots that show the accuracy of the estimated selection coefficients in the respective scenarios. If a method was not applicable in a given scenario, we omitted it from the boxplot. We observe that in the scenarios that a certain method is designed for, the coefficients are estimated fairly accurately. `diplo-locus` achieves the best accuracy in all scenarios or is at least tied with the best method. All other methods do better in some scenarios, but worse in others. Note that in the case of dominance, the estimates that were produced using `LLS` are accurate, however, the method produces estimates of `nan` for 19% of

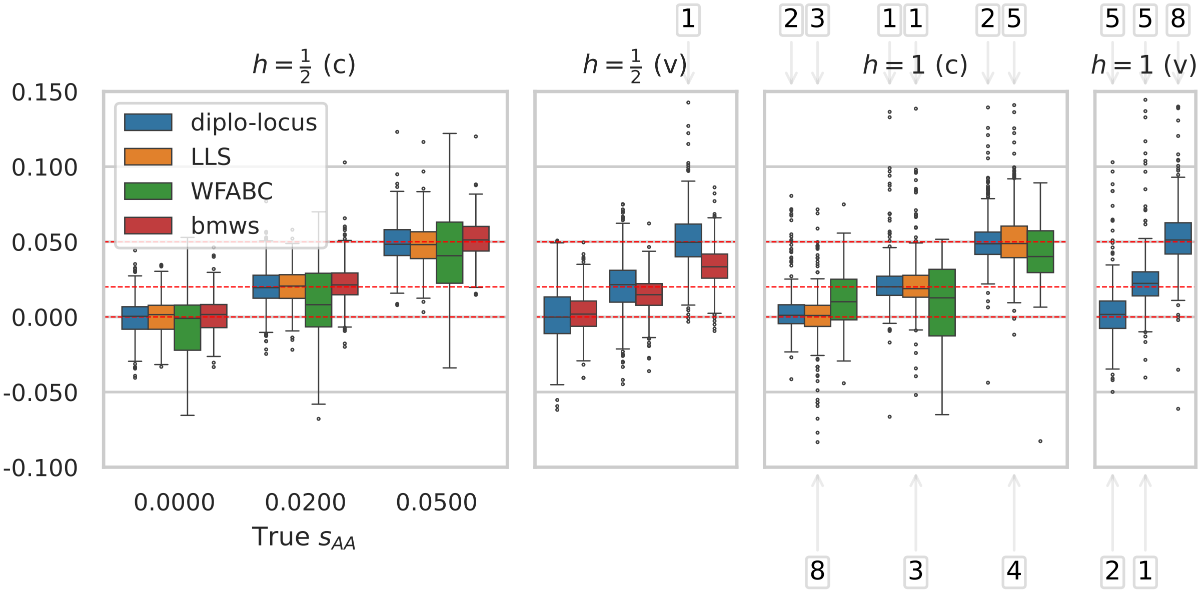

Figure S14: Boxplots exhibiting inference accuracy of the methods `diplo-locus`, `LLS`, `WFABC`, and `bmws`. Simulations for 160 generations, initialized from standing variation, sampled at 9 timepoints. The scenarios are either additive selection ($h = \frac{1}{2}$) or dominance ($h = 1$), and either a single constant coefficient (c) or a scenario where the coefficient changes (v). We only show results for $s \in \{0, 0.02, 0.05\}$, and omit a method, if it was not designed for a certain scenario. Numbers above and below plot denote estimates out of plot range. Only 500 out of 2000 replicates shown.

the replicates when simulating for 160 generations and for 77% of the replicates when simulating for 4000 generations. The method `bmws` slightly underestimates the selection coefficients in the 160 generation scenario with time-varying selection. This is likely due to the fact that the method was designed for gradual changes in the selection coefficient. In addition, in the scenarios with 4000 generations, `bmws` estimates $s = 0$ for all replicates, which is likely because the regularization parameter recommended by the authors is not well-calibrated here. However, we were not able to identify a regularization parameter that led to non-zero estimates. Overall, this comparison shows that `diplo-locus` can be used for accurate inference in a variety of scenarios.

### S5 Analyses of Empirical Data

#### S5.1 Log-likelihood surface for *ASIP* locus in ancient horses

We use the temporal allele counts at the *ASIP* locus in the dataset obtained from ancient horses by Ludwig et al. (2009) as presented by Steinrücken et al. (2014), see Table S2. We assume equal forward and backward mutation rates, $u_{Aa}$ and $u_{aA}$, to be $10^{-6}$ per site per generation, with a generation time of 8 years. Following He et al. (2023), we assume the effective population size to

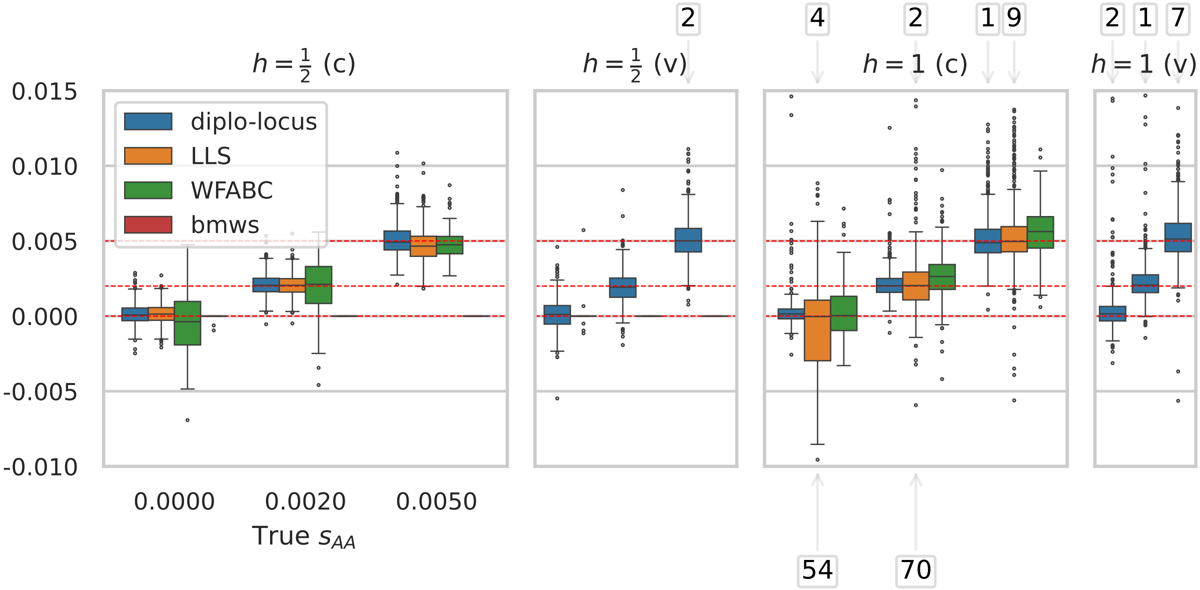

Figure S15: Boxplots exhibiting inference accuracy of the methods `diplo-locus`, `LLS`, `WFABC`, and `bmws`. Simulations for 4000 generations, initialized from standing variation, sampled at 9 timepoints. The scenarios are either additive selection ($h = \frac{1}{2}$) or dominance ($h = 1$), and either a single constant coefficient (c) or a scenario where the coefficient changes (v). We only show results for $s \in \{0, 0.002, 0.005\}$, and omit a method, if it was not designed for a certain scenario. Numbers above and below plot denote estimates out of plot range. Only 500 out of 2000 replicates shown.

Table S2: Allele count data at the *ASIP* locus in ancient horses used for the analyses. The mutation is assumed to occur at $t_0 = 17,000$ years before the common era (BCE) with an initial frequency of $1/(2N_e)$.

| Sampling time (BCE) | 20,000 | 13,100 | 3,700 | 2,800 | 1,100 | 500 |
|---|---|---|---|---|---|---|
| Total number of alleles observed ($n$) | 10 | 22 | 20 | 20 | 36 | 38 |
| Number of derived alleles ($d$) | 0 | 1 | 15 | 12 | 15 | 18 |

be constant in the relevant time-frame at $N_e = 16,000$, consistent with the estimates obtained by Der Sarkissian et al. (2015). To compute the log-likelihoods, we assume that the favored allele arose from a *de novo* mutation, that is, its initial frequency is $1/(2N_e)$, and it is subject to constant selective pressure.

We computed log-likelihoods for three different possible times when the selected mutation arose $t_0 \in \{17000, 15000, 13105\}$ years before the common era (BCE), where the most recent time is immediately before the allele is first observed in the sample. We computed log-likelihoods on a two-dimensional grid for $s_{\text{aA}}$ and $s_{\text{AA}}$ values in $(-0.1, 0.1)$ with a step size of 0.004. The log-likelihood surfaces are depicted in Figure S16. We observe that the maximum likelihood estimate for the selection coefficients is consistent with balancing selection, as previously reported by Steinrücken et al. (2014), and is not strongly affected by different assumed times of origin of the selected allele. The most recent time $t_0 = 13,105$ yields the highest log-likelihood, but the difference is not large, indicating that there is not much power to estimate the time of origin from the given dataset. We also depict the surface for $t_0 = 13,105$ in Figure 1D) in the main text. Scripts for the analysis can be found at `https://github.com/steinrue/diplo_locus_manuscript_figs`, but are also included as Example 1 in the tutorial section of the documentation for our software (`https://github.com/steinrue/diplo_locus`).

##### S5.2 Analysis of temporal allele counts of ancient humans in Great Britain

Following a similar rationale as Mathieson and Terhorst (2022), we extracted 520 unrelated ancient human genomes sampled in Great Britain 4,500 years before present (1950 CE) from the Allen Ancient DNA Resource 1240K v54.1 dataset (Mallick et al., 2024). We only included unrelated samples annotated as "PASS" and assessed on the 1240K capture panel. Note that this excluded all contemporary samples. The reason to only analyze capture data is to avoid potential confounding due to analyzing capture and Shotgun data jointly (Margaryan et al., 2020, Suppl. Note 8). This effectively resulted in a subset of the samples published by Schiffels et al. (2016), Martiniano et al. (2016), Olalde et al. (2018), Patterson et al. (2022), and Gretzinger et al. (2022).

Focusing on Chromosome 2, we extracted pseudo-haploid genotypes at 98,657 SNPs on this chromosome for the given individuals. Grouping the samples into generations, using a generation time of 28.1 years (Moorjani et al., 2016), these SNPs have been recorded at 94 time points, with sample sizes varying from 1 to 52. We removed SNPs whose pooled MAF is below 0.05, where fewer than 6 samples (around 1% of the total number of samples) have variant calls, or variant calls exist at fewer than three time points. This resulted in a dataset with 69,901 SNPs after filtering.

We used the command line tool `DiploLocus likelihood` to compute log-likelihoods at these filtered SNPs across Chromosome 2, assuming a mutation rate of $1.25 \times 10^{-8}$ per-generation per-base, a uniform initial condition and constant additive selection ($h = 0.5$). Moreover, we specify a

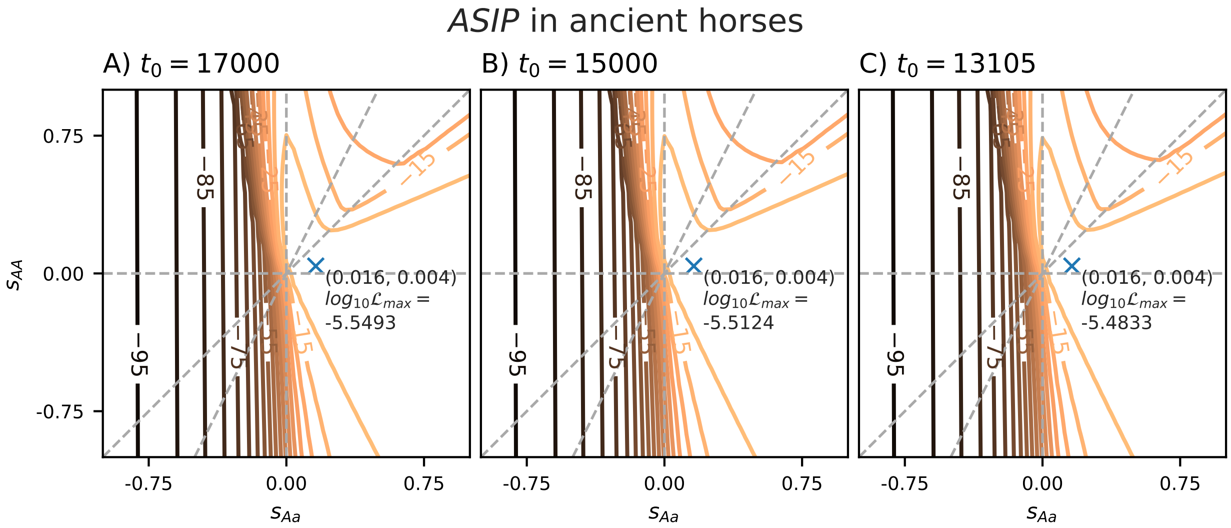

Figure S16: Log-likelihood surfaces computed from the temporal allele frequency data at the ASIP locus in ancient horses. We vary the assumed time of origin of the selected allele between A) $t_0 = 17,000$, B) $t_0 = 15,000$, and C) $t_0 = 13,105$, where the latter time yields the highest log-likelihood at the maximum.

single constant population size for the time period of sampling. To obtain this value, we use the estimates provided by (Browning et al., 2015, Figure 4A), who estimated variable past effective population sizes in Great Britain using shared haplotype information in the UK10K dataset. Our most ancient sample is dated 4480 years before present and our most recent sample is dated 856 years before present. From Figure 4A by Browning et al. (2015) we can gauge that the effective population size increased from approximately $10^4$ to approximately $4 \cdot 10^5$ during this period. Assuming exponential growth, this corresponds to approximately 3% growth per generation. Under the assumption of growth at this rate, we take the harmonic mean of the effective population sizes, resulting in approximately 37,000, which we then use as the effective population size $N_e$ for the analysis with `DiploLocus likelihood`.

We computed log-likelihoods for each SNP along a one-dimensional symmetric geometric grid of $s_{\mathrm{AA}}$ values in the interval [-0.5, 0.5] (51 values in total; zero included), interpolate to obtain off-grid MLEs and the corresponding LLRs, and calculated $p$-values using a $\chi^2(1)$ distribution. Figure S17 (see also Figure 2b in the main text) shows a Manhattan plot of all $p$-values on Chromosome 2, and Figure S18 shows the $p$-values in the 2 Mbp genomic region around the lactase gene *LCT*. Additionally, we show a log-likelihood surface varying both selection coefficients in Figure 2c in the main text.

The most significant SNP is `rs4988235`, which is located in an intron of the *MCM6* gene and upstream of the *LCT* gene, and other significant SNPs cluster around this SNP. No other SNPs on Chromosome 2 besides this cluster pass a Bonferroni-corrected threshold. The signal of selection in this genomic region has been well characterized in many studies and has a well-supported connection to lactase persistence (Enattah et al., 2002; Troelsen et al., 2003). Scripts for the analysis can be found at `https://github.com/steinrue/diplo_locus_manuscript_figs`, but are also included as Example 2 in the tutorial section of the documentation for our software

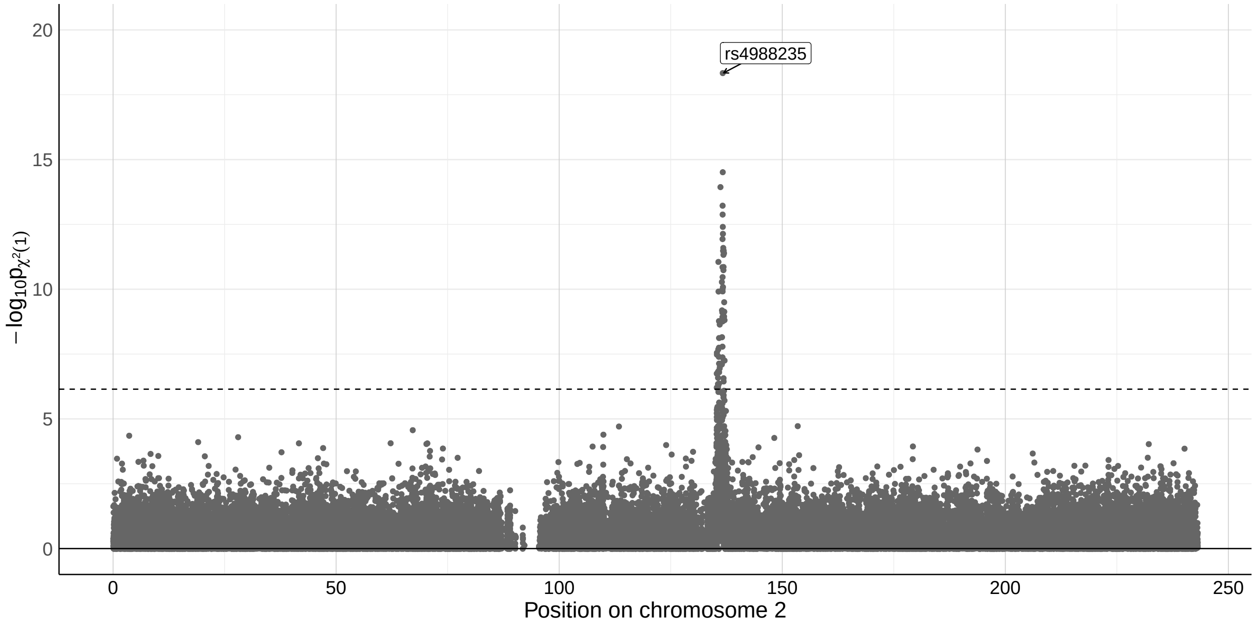

Figure S17: Manhattan plot of $p$-values for filtered SNPs on Chromosome 2 in temporally-stratified samples from Great Britain extracted from the Allen ancient DNA Resource Database (AADR, v54.1). Likelihood computation assumes a uniform initial distribution, additive selection, and uses a symmetric geometric grid covering [-0.5, 0.5] for $s_{\mathrm{AA}}$ with 51 points. Dashed line indicates the negative logarithm of the Bonferroni-corrected threshold at $p = 0.05$.

(`https://github.com/steinrue/diplo_locus`).

##### S5.3 Analysis of *Drosophila simulans* Evolve & Resequence dataset

We used `diplo-locus` to analyze an Evolve & Resequence dataset collected by Barghi et al. (2019). In this experiment, the authors exposed 10 biologogical replicates of *Drosophile simulans* populations to a new temperature regime. The replicate populations were initialized using 202 isofemale lines, and the allele frequencies in the population where assessed every 10 generations, from generation 0 until generation 60, using pooled-sequencing.

For each replicate, and at each timepoint, we extracted the number of sequencing reads for the major and the minor allele at each of the 1,068,826 biallelic loci on chromosome 2L. We furthermore applied a filter for the minor allele frequency (MAF) by pooling the samples across timepoints at each locus, and discarding loci where the MAF was below 0.05. This resulted in approximately 700,000 biallelic loci per biological replicate.

We then applied `diplo-locus` to this temporal allele frequency data at each locus, considering the sequencing reads as binomial samples conditional on the underlying population allele frequency, the emission model of `diplo-locus`. At each locus, we computed the likelihoods under a model of additive selection, for values of the coefficient $s$ on a geometric grid of 63 points in $[-0.75, 0.75]$ (including 0). We used the uniform distribution for the initial frequency, a mutation rate of $5 \cdot 10^{-9}$ per generation, and $N_e = 300$, consistent with Table S5 given by Barghi et al. (2019). We then used these likelihoods to get an MLE of the additive selection coefficient at each locus, and applied the $\chi^2$

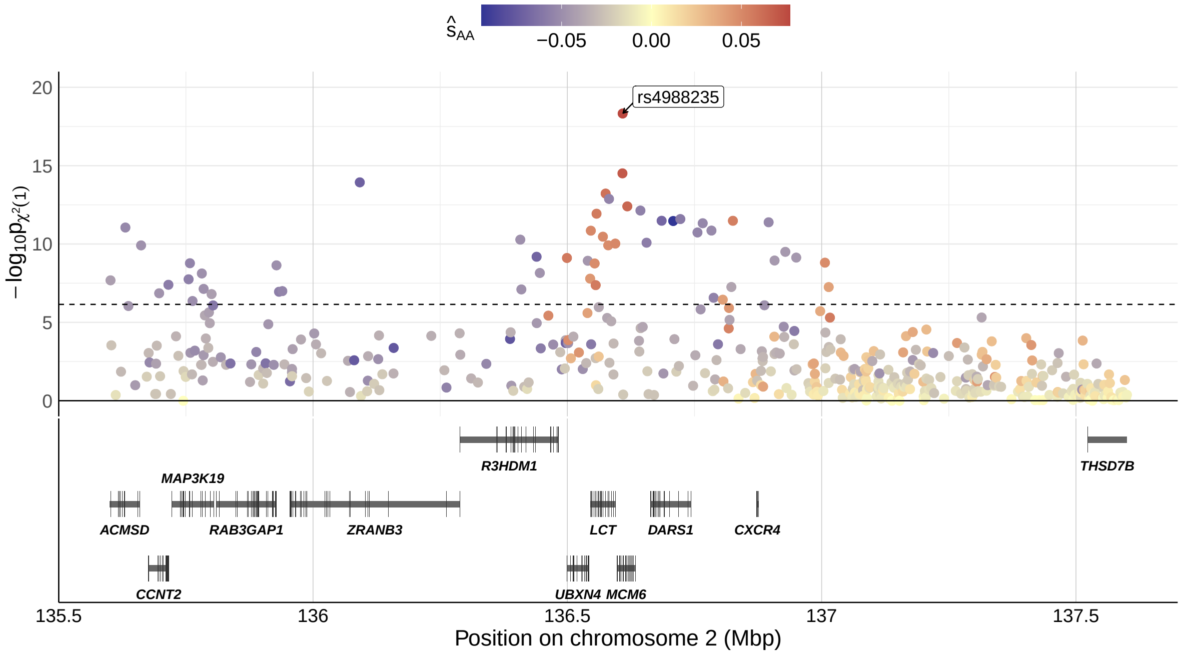

Figure S18: Manhattan plot of $p$-values for additive selection in the 2 Mbp region approximately centering on the *LCT*/*MCM6* locus. Color codes for the value of the MLEs for $s_{\mathrm{AA}}$. The longest transcripts of protein-coding genes are shown below the Manhattan plot.

distribution with 1 degree of freedom to the likelihood ratio statistic at each locus to obtain $p$-values assessing the evidence for a non-zero selection coefficient. Since the data comprised 10 biological replicates of the experiment, this resulted in 10 $p$-values at each locus, one for each experiment. To asses the combined evidence for a non-zero selection coefficient across the 10 biological replicates, we applied Fisher's method (Mosteller and Fisher, 1948) to the subset of common loci, resulting in one $p$-value per locus. Figure 1a in the main text and Figure S19 show a Manhattan plot of the $p$-values, as well as a threshold obtained from Bonferroni correction for multiple testing at a significance level of 0.05.

While a detailed analysis of the results in the context of the experiment is beyond the scope of this manuscript, Figure S20 and Figure S21 show the $p$-values around the two clearly visible peaks. The names of the genes were obtained using the FlyBase ID annotations from Palmieri et al. (2015) of orthologs in Drosophila melanogaster to query the database at `https://www.alliancegenome.org/`. Based on the information in this database, the *CG*-genes and *E23* in Figure S20 are related to transmembrane transporter activity. The *CG*-genes in Figure S21 have no annotated function, but *Ddr* is involved in protein tyrosine kinase signaling, and the human ortholog is implicated in several diseases.

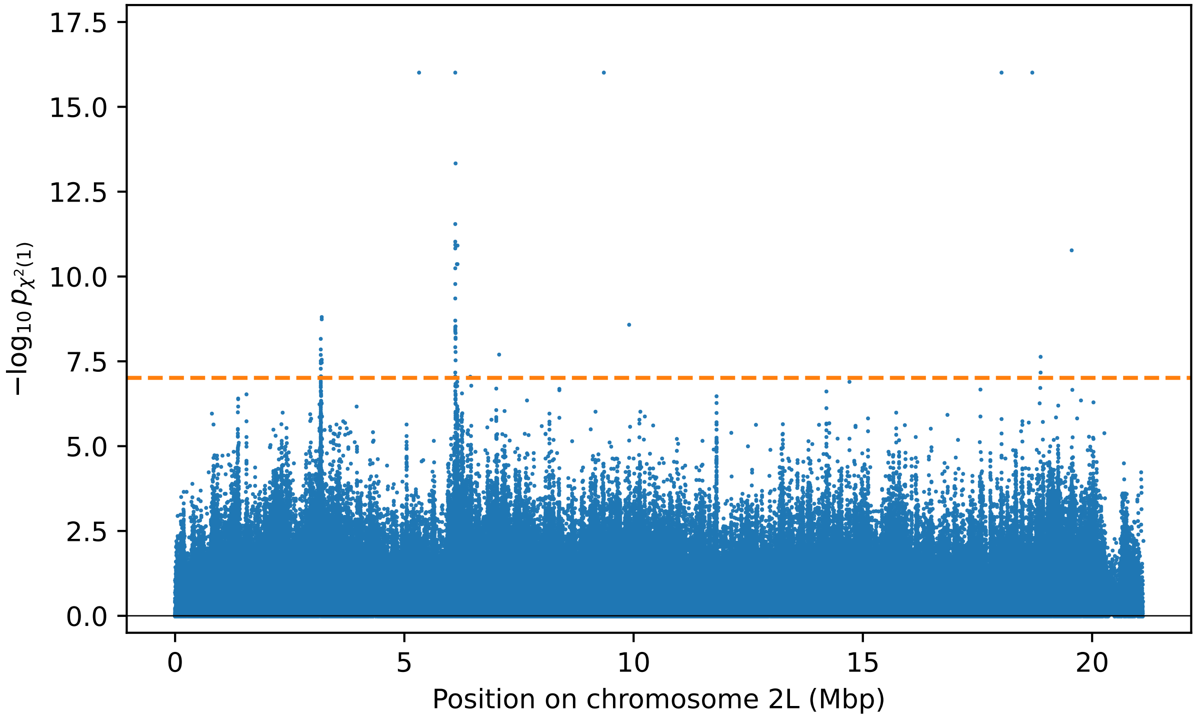

Figure S19: Manhattan plot of $p$-values for additive selection on chromosome 2L. Horizontal dashed line shows Bonferroni correction at level 0.05.

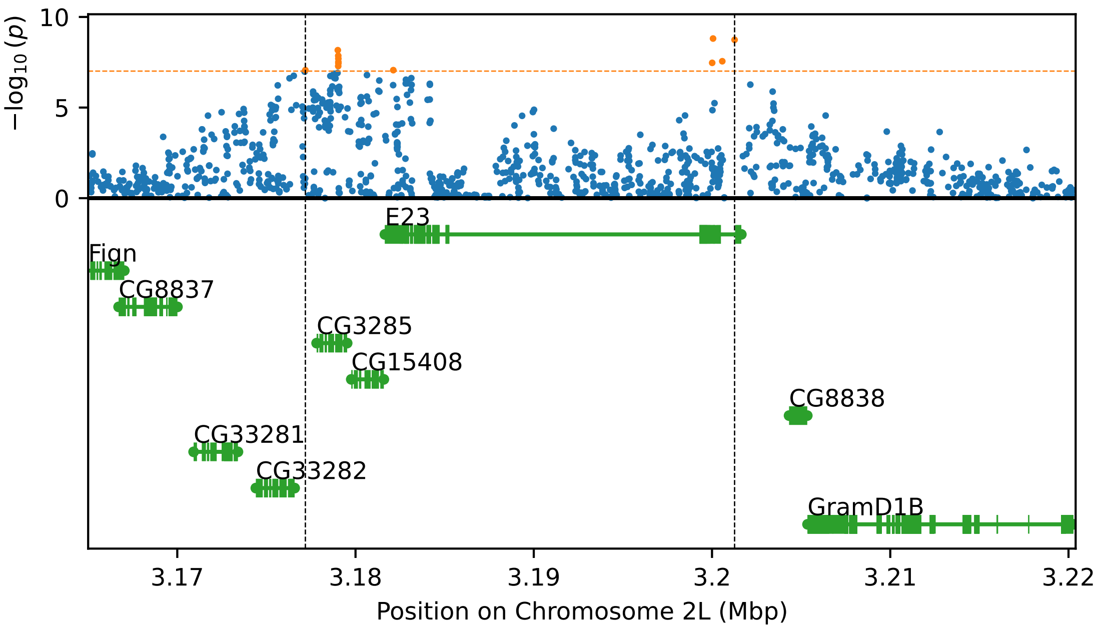

Figure S20: Manhattan plot of $p$-values around first strong peak on chromosome 2L and genes in the genomic region. Horizontal dashed line shows Bonferroni correction at level 0.05.

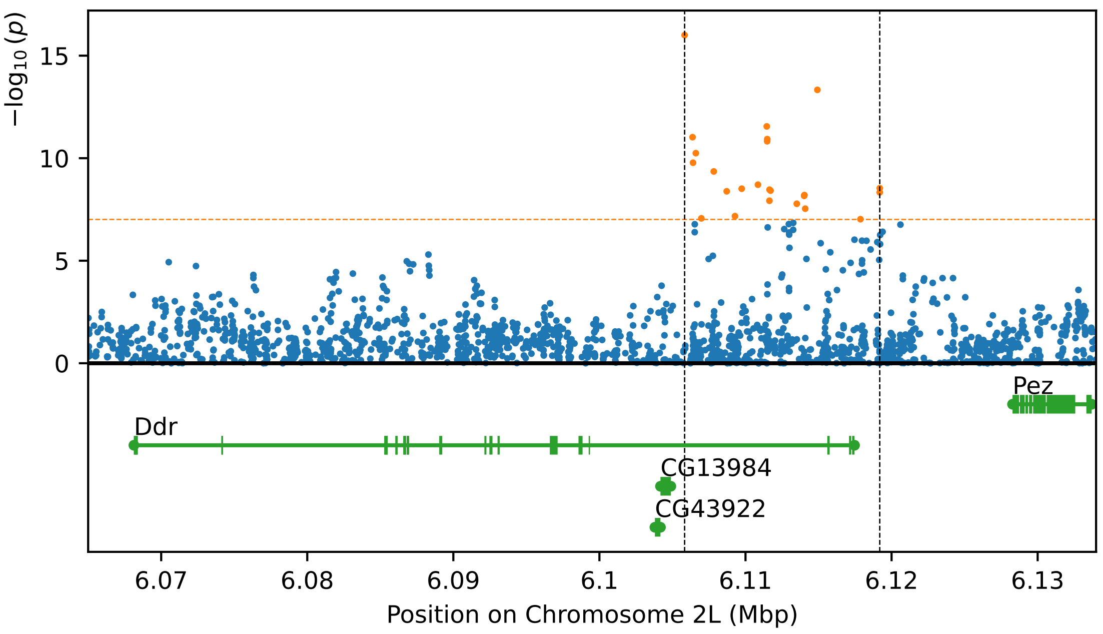

Figure S21: Manhattan plot of $p$-values around second strong peak on chromosome 2L and genes in the genomic region. Horizontal dashed line shows Bonferroni correction at level 0.05.

Baumdicker, F., Bisschop, G., Goldstein, D., Gower, G., Ragsdale, A. P., Tsambos, G., Zhu, S., Eldon, B., Ellerman, E. C., Galloway, J. G., et al. 2022. Efficient ancestry and mutation simulation with msprime 1.0. *Genetics*, 220(3):iyab229.

Bishop, C. M. *Pattern Recognition and Machine Learning.* Springer, 2016.

Bollback, J. P., York, T. L., and Nielsen, R. 2008. Estimation of $2N_e s$ from temporal allele frequency data. *Genetics*, 179(1):497–502.

Browning, M., Behrens, T. E., Jocham, G., O'Reilly, J. X., and Bishop, S. J. 2015. Anxious individuals have difficulty learning the causal statistics of aversive environments. *Nature Neuroscience*, 18(4):590–596.

Cheng, X. and DeGiorgio, M. 2020. Flexible mixture model approaches that accommodate footprint size variability for robust detection of balancing selection. *Molecular Biology and Evolution*, 37: 3267–3291.

Der Sarkissian, C., Ermini, L., Schubert, M., Yang, M. A., Librado, P., Fumagalli, M., Jónsson, H., Bar-Gal, G. K., Albrechtsen, A., Vieira, F. G., Petersen, B., Ginolhac, A., Seguin-Orlando, A., Magnussen, K., Fages, A., Gamba, C., Lorente-Galdos, B., Polani, S., Steiner, C., Neuditschko, M., Jagannathan, V., Feh, C., Greenblatt, C. L., Ludwig, A., Abramson, N. I., Zimmermann, W., Schafberg, R., Tikhonov, A., Sicheritz-Ponten, T., Willerslev, E., Marques-Bonet, T., Ryder, O. A., McCue, M., Rieder, S., Leeb, T., Slatkin, M., and Orlando, L. 2015. Evolutionary genomics and conservation of the endangered przewalski's horse. *Current Biology*, 25(19):2577–2583.

Durrett, R. *Probability models for DNA sequence evolution.* Springer, 2008.

Enattah, N. S., Sahi, T., Savilahti, E., Terwilliger, J. D., Peltonen, L., and Järvelä, I. 2002. Identification of a variant associated with adult-type hypolactasia. *Nature Genetics*, 30(2):233–237.

Foll, M., Shim, H., and Jensen, J. D. 2015. WFABC: a wright-fisher abc-based approach for inferring effective population sizes and selection coefficients from time-sampled data. *Molecular Ecology Resources*, 15(1):87–98.

Gretzinger, J., Sayer, D., Justeau, P., Altena, E., Pala, M., Dulias, K., Edwards, C. J., Jodoin, S., Lacher, L., Sabin, S., et al. 2022. The Anglo-Saxon migration and the formation of the early English gene pool. *Nature*, 610(7930):112–119.

Haller, B. C. and Messer, P. W. 2023. SLiM 4: multispecies eco-evolutionary modeling. *The American Naturalist*, 201(5):E000–E000.

He, Z., Dai, X., Lyu, W., Beaumont, M., and Yu, F. 2023. Estimating temporally variable selection intensity from ancient dna data. *Molecular Biology and Evolution*, 40(3):msad008.

Ludwig, A., Pruvost, M., Reissmann, M., Benecke, N., Brockmann, G. A., Castaños, P., Cieslak, M., Lippold, S., Llorente, L., Malaspinas, A.-S., et al. 2009. Coat color variation at the beginning of horse domestication. *Science*, 324(5926):485–485.

Mallick, S., Micco, A., Mah, M., Ringbauer, H., Lazaridis, I., Olalde, I., Patterson, N., and Reich, D. 2024. The allen ancient dna resource (AADR) a curated compendium of ancient human genomes. *Scientific Data*, 11(1):182.

Margaryan, A., Lawson, D. J., Sikora, M., Racimo, F., Rasmussen, S., Moltke, I., Cassidy, L. M., Jørsboe, E., Ingason, A., Pedersen, M. W., et al. 2020. Population genomics of the Viking world. *Nature*, 585(7825):390–396.

Martiniano, R., Caffell, A., Holst, M., Hunter-Mann, K., Montgomery, J., Müldner, G., McLaughlin, R. L., Teasdale, M. D., Van Rheenen, W., Veldink, J. H., et al. 2016. Genomic signals of migration and continuity in Britain before the Anglo-Saxons. *Nature communications*, 7(1): 10326.

Mathieson, I. and McVean, G. 2013. Estimating selection coefficients in spatially structured populations from time series data of allele frequencies. *Genetics*, 193(3):973–984.

Mathieson, I. and Terhorst, J. 2022. Direct detection of natural selection in Bronze Age Britain. *Genome Research*, 32(11-12):2057–2067.

Moorjani, P., Sankararaman, S., Fu, Q., Przeworski, M., Patterson, N., and Reich, D. 2016. A genetic method for dating ancient genomes provides a direct estimate of human generation interval in the last 45,000 years. *Proceedings of the National Academy of Sciences*, 113(20): 5652–5657.

Mosteller, F. and Fisher, R. A. 1948. Questions and answers #14. *The American Statistician*, 2 (5):30–31.

Olalde, I., Brace, S., Allentoft, M. E., Armit, I., Kristiansen, K., Booth, T., Rohland, N., Mallick, S., Szécsényi-Nagy, A., Mittnik, A., et al. 2018. The beaker phenomenon and the genomic transformation of northwest europe. *Nature*, 555(7695):190–196.

Palmieri, N., Nolte, V., Chen, J., and Schlötterer, C. 2015. Genome assembly and annotation of a drosophila simulans strain from madagascar. *Molecular Ecology Resources*, 15(2):372–381.

Patterson, N., Isakov, M., Booth, T., Büster, L., Fischer, C.-E., Olalde, I., Ringbauer, H., Akbari, A., Cheronet, O., Bleasdale, M., et al. 2022. Large-scale migration into Britain during the Middle to Late Bronze Age. *Nature*, 601(7894):588–594.

Sawyer, S. A. and Hartl, D. L. 1992. Population genetics of polymorphism and divergence. *Genetics*, 132(4):1161–1176.

Schiffels, S., Haak, W., Paajanen, P., Llamas, B., Popescu, E., Loe, L., Clarke, R., Lyons, A., Mortimer, R., Sayer, D., et al. 2016. Iron age and Anglo-Saxon genomes from East England reveal British migration history. *Nature communications*, 7(1):10408.

Self, S. G. and Liang, K.-Y. 1987. Asymptotic properties of maximum likelihood estimators and likelihood ratio tests under nonstandard conditions. *Journal of the American Statistical Association*, 82(398):605–610.

Steinrücken, M., Bhaskar, A., and Song, Y. S. 2014. A novel spectral method for inferring general diploid selection from time series genetic data. *The Annals of Applied Statistics*, 8(4):2203–2222.

Taus, T., Futschik, A., and Schlötterer, C. 2017. Quantifying selection with pool-seq time series data. *Molecular Biology and Evolution*, 34(11):3023–3034.

Troelsen, J. T., Olsen, J., Møller, J., and Sjöström, H. 2003. An Upstream Polymorphism Associated with Lactase Persistence has Increased Enhancer Activity. *Gastroenterology*, 125(6): 1686–1694.

Watterson, G. A. 1975. On the number of segregating sites in genetical models without recombination. *Theoretical Population Biology*, 7(2):256–276.
